# Motor implementation of control and reward-based urgency regulation across impulsivity

**DOI:** 10.64898/2026.01.30.702817

**Authors:** Thibault Fumery, Fostine Chaise, Anaelle Soille Hambye, Fanny Fievez, Julien Lambert, Pierre Vassiliadis, Gerard Derosiere, Julie Duque

## Abstract

Everyday decisions unfold dynamically, with commitment shaped by a growing sense of urgency that can, when excessive, contribute to impulsive choices. Here we aimed at dissociating two modes of urgency regulation, control-driven (accuracy-oriented) and reward-driven (motivation-based), and asked whether their relative influence varies across individuals differing in impulsivity. We further investigated how these regulatory modes are implemented in the motor system, focusing on two modulatory effects: surround inhibition and broad modulation. Healthy participants, whose impulsivity was assessed with the UPPS urgency dimension, performed a modified Tokens task crossing control demands (low vs high control blocks) with motivational context (low vs high reward trials). In two separate sessions, single-pulse TMS was applied either over the hand motor representation to probe corticospinal excitability indexing surround inhibition, or over the leg representation to index broad modulations of motor activity. This design successfully dissociated the two regulatory modes: control-driven adjustments (across blocks) were most evident in less impulsive participants, whereas reward-driven adjustments (across trials) were most evident in more impulsive participants. Consistent with this dissociation, control-driven urgency regulation was associated with broad modulation of motor activity, whereas reward-driven urgency adjustments were associated with changes in surround inhibition. These motor signatures may serve as probes of the respective contributions of control- and reward-driven regulation even when they are not explicitly dissociated. Our findings suggest that impulsivity may not simply reflect “more urgency” but a different weighting of the influences that shape it during decision making, a hypothesis that can now be tested in clinical conditions.

## 1. Introduction

Many everyday decisions are dynamic and action–based, unfolding “on the fly” as situations evolve. For example, while moving through the aisle of a wine shop to pick a bottle before a dinner party, your preference gradually changes as new cues appear (e.g., labels, prices, producers) until you ultimately commit to a single option. Growing work suggests that these choices are shaped by an urgency signal that rises from deliberation onset and drives commitment (Yau et al., 2021; Thura et al., 2022). Accordingly, urgency has been proposed as a substrate for impulsivity (Carland et al., 2019), a tendency toward premature commitment; however, whether urgency tracks inter-individual variation in impulsivity, from typical population differences to extreme psychopathology, remains unknown.

Urgency is not fixed within an individual but can be regulated in a goal-directed manner. When accuracy is prioritized, urgency is typically reduced, consistent with top-down engagement of inhibitory control to delay commitment and allow further evidence sampling (Thura, 2020; Derosiere et al., 2021; Fievez et al., 2024). In the wine-shop example, being instructed to bring a “good” bottle prioritizes accuracy, encouraging additional sampling and the exertion of control over the urge to join the party. This effect is even stronger when the outcome is more rewarding (e.g., for a wine enthusiast), such that motivation can further amplify this accuracy-oriented control-driven caution and temper urgency (Manohar et al., 2015; Frömer et al., 2021; Reppert et al., 2023). Here, we aimed to dissociate such control-driven and reward-driven modes of urgency regulation. Moreover, building on evidence that impulsivity is associated with reduced ability to engage inhibitory control but heightened reward sensitivity (Drew et al., 2020; Barakat et al., 2025), we hypothesized that urgency would be largely control-regulated in less-impulsive individuals, whereas more impulsive individuals would show stronger reward-based modulation..

Action-based decisions recruit modulatory processes in the motor system (Thura et al., 2025), yet how urgency shapes these motor dynamics remains unclear. Prior work suggests the operation of two complementary adjustments consistent with the view that urgency is jointly regulated by control-driven and reward-driven influences. First, prior studies have linked urgency changes to broad, non-selective shifts in motor excitability (Steinemann et al., 2018; Murphy et al., 2020; Derosiere et al., 2022), interpreted as reflecting top-down engagement of inhibitory control that tunes the effective distance to commitment: stronger inhibition increases this distance under accuracy goals, whereas weaker inhibition reduces it under speed goals, facilitating faster commitment. Building on this interpretation, we treat broad modulation as the candidate motor implementation of control-driven urgency adjustments. Second, urgency regulation has been related to more selective, local inhibitory mechanisms such as surround inhibition, which sharpens the selected effector’s activity by suppressing nearby, irrelevant muscles (Leodori et al., 2019; Derosiere et al., 2022). Consistently, prior work has linked surround inhibition to motor gain control and reward-related neuromodulatory influences (Derosiere et al., 2025), suggesting that this selective inhibitory mechanism could be a route through which reward-driven motivation shapes urgency. Accordingly, we consider changes in surround inhibition as the candidate motor signature of reward-driven urgency regulation. Moreover, consistent with our behavioral hypothesis and its link to impulsivity, we predict that the relative expression of these motor signatures will vary with the impulsivity level: control-driven broad modulation will be most evident in less impulsive individuals, whereas reward-driven changes in surround inhibition will be most evident in more impulsive subjects.

Here, we aimed to dissociate control-driven and reward-driven modes of urgency regulation and to examine their motor implementation across healthy individuals differing in impulsivity. To this end, we adapted the Tokens task to vary the required control level (low vs. high control) and motivational context (low vs. high reward), predicting that these two manipulations would differentially recruit these two regulatory modes. Under this framework, we aimed to identify the motor signatures through which these urgency adjustments are implemented in the motor system across the impulsivity spectrum in the general population.

## 2. Materials and Methods

### 2.1 Participants, Ethics and Questionnaires

#### Participants

A total of twenty-five healthy individuals were recruited, of whom twenty were included in the final analyses (13 women, 24.2 ± 3.2 years), as explained later in the methods (see Transcranial Magnetic Stimulation section). They were all right-handed, as assessed by the Edinburgh Handedness Inventory (Oldfield, 1971) (i.e., laterality score above 80%), and had normal or corrected-to-normal vision. Exclusion criteria included any neurological or psychiatric disorder, any drug treatment that could influence performance or neural activity, or any history of substance use disorder.

#### Ethics statement

The protocol was approved by the Ethics Committee of the Université catholique de Louvain, Brussels, Belgium (approval number: AAHR-DQS-037), and adhered to the Declaration of Helsinki. All participants provided written informed consent and received financial compensation for their participation.

#### Questionnaires

Participants first completed a medical screening questionnaire to confirm their eligibility and rule out any medical contraindications to brain stimulation. They then completed the UPPS-P Impulsive Behavior Scale (short 20-item French version) (Cyders et al., 2014), a self-report questionnaire assessing trait impulsivity. This scale measures four distinct dimensions of impulsivity: urgency, lack of premeditation, sensation seeking and lack of perseverance. Each dimension is quantified by averaging item responses scored from 1 to 4, in which higher values reflect greater trait impulsivity. Here, we focused on the urgency subscale (UPPS_Urgency,_ including both positive and negative urgency), which emerged as the dimension showing the most robust and reliable effects in our analyses.

### 2.2 Experimental setup and task

The experiment was conducted in a quiet and dimly lit room. Participants were comfortably seated in front of a 21-inch cathode ray tube monitor, positioned at approximately 60 cm from their eyes, and used to display stimuli during the task. Participants’ forearms were positioned in a semi-pronated position on wooden plates placed on the table. To ensure a fixed and standardized resting position, elastic bands were used to hold each index finger in a full extension position in between trials of the task. To minimize external auditory distractions, participants wore earphones delivering continuous white noise. The task used in the present study is a modified version of the Tokens task (Cisek et al., 2009; Thura & Cisek, 2014), implemented in Matlab (The Mathworks, Natick, Massachusetts, USA) with the Cogent 2000 toolbox (Functional Imaging Laboratory, London, UK).

#### General principle of the Tokens task

The Tokens task required participants to decide which of two lateral circles would receive the majority of 15 tokens jumping progressively from a central circle. As shown on Figure 1A (upper part of left panel), all 15 tokens were initially distributed at random positions in the central circle. One token then jumped to a lateral circle, and the remaining tokens followed one by one at 200-ms intervals, each jumping to either the left or right circle. Participants had to predict which lateral circle would ultimately contain the majority of tokens and indicate their choice by performing three consecutive flexions of the same left or right index finger, corresponding to the left or right circle, respectively. Two lasers targeted the fingertip of each index to define their initial resting position (i.e., 0° flexion) and the movement amplitude required by the tapping movement (i.e., 45° flexion, see Fig. 1A, lower part of left panel). Participants were instructed to respond once they felt sufficiently confident, but always before the last (15^th^) token jump (Jump_15_). After the finger response, the remaining tokens continued to jump at the same pace until the central circle was empty.

**Figure 1.**
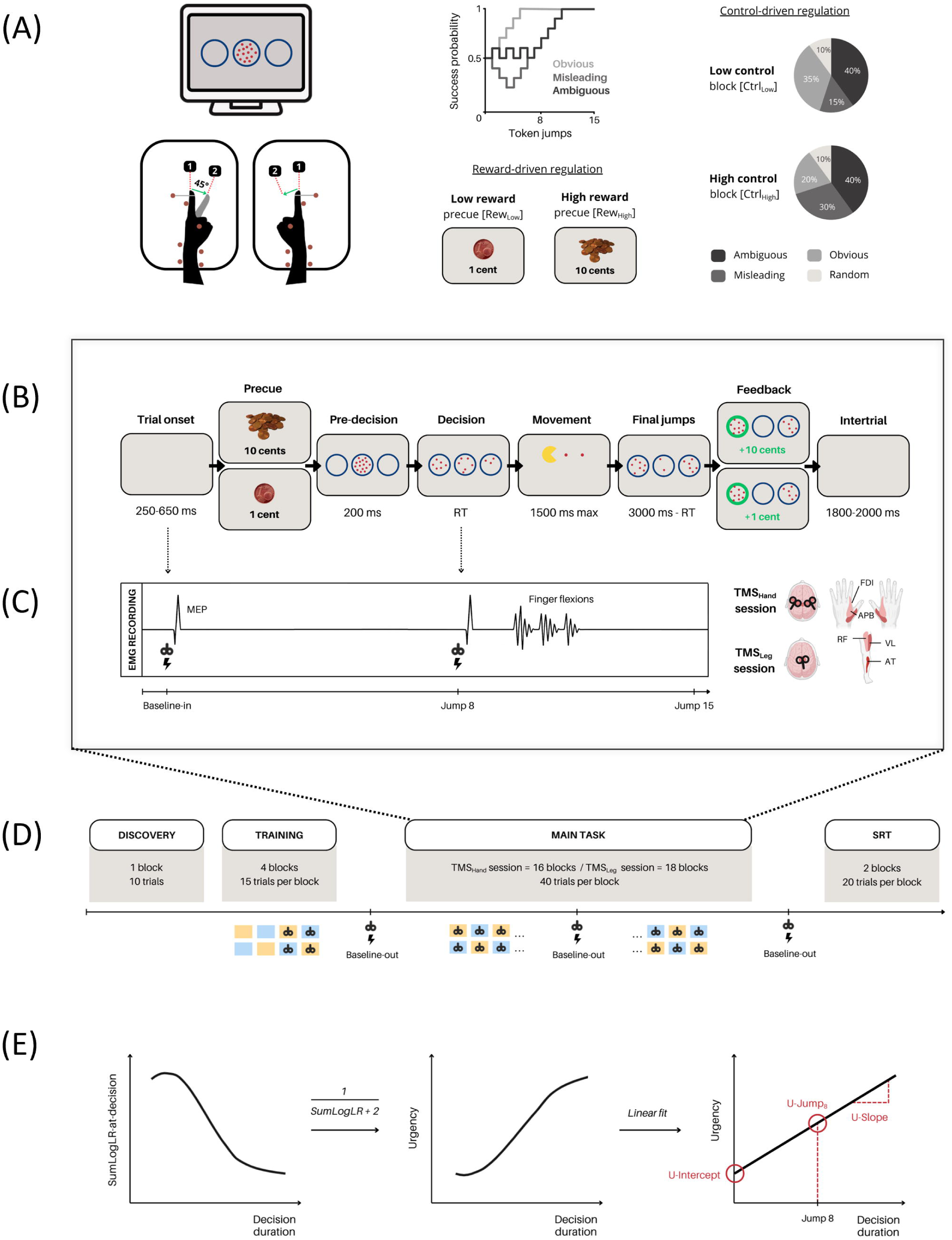
Experimental design, TMS protocol, and urgency modelling. **(A) Tokens task and experimental conditions. Left panel:** *Upper part:* At trial onset, three circles appeared at the center of the screen, with 15 tokens initially displayed in the central circle. *Lower part:* Participants’ forearms rested on wooden plates, with small wooden sticks secured around each forearm to ensure consistent placement. Elastic bands maintained both index fingers in the starting position. Two lasers (targeting the tip of each index finger) indicated the starting position (0° flexion; laser 1) and the required movement amplitude (45° flexion; laser 2). **Middle panel:** *Upper part:* The task involved different trial types, defined by how success probability evolved over time. In obvious trials (light grey), success probability rose quickly to high values; in misleading trials (medium grey), it was initially low before increasing later; and in ambiguous trials (dark grey), it remained intermediate for longer before eventually increasing. *Lower part:* To promote reward-driven adjustments of urgency, the task included two precue types: Rew_Low_ signaled a low reward (1 cent) and Rew_High_ signaled a high reward (10 cents) for successful trials, each occurring on half of the trials in a random order. **Right panel:** To promote control-driven adjustments of urgency, the task included two block types: Ctrl_Low_ blocks contained more obvious trials (35%) and fewer misleading trials (15%), whereas Ctrl_High_ blocks contained fewer obvious trials (20%) and more misleading trials (30%). **(B) Trial sequence of events.** Each trial comprised a reward precue, a pre-decision display of the three circles with tokens, a decision period during which tokens jumped to the lateral circles, and a movement report (three left/right index-finger flexions; Pacman animation). Final jumps then continued until all tokens had moved, followed by accuracy feedback and a variable intertrial interval. **(C) Transcranial Magnetic Stimulation (TMS) procedure. Left panel:** In the TMS_Hand_ session, a double-coil approach was used to stimulate the primary motor cortex (M1) bilaterally to elicit Motor-evoked potentials (MEPs) in both hands, in the First Dorsal Interosseous (FDI) and in the Abductor Pollicis Brevis (APB) muscles. In the TMS_Leg_ session, TMS was applied on right M1 to elicit MEPs in 3 muscles of the left leg: the Vastus Lateralis (VL), the Rectus Femoris (RF) and the Tibialis Anterior (TA). **Right panel:** In both sessions, TMS pulses were delivered at two time points within task blocks: at baseline (TMS_Baseline-in_), during the intertrial interval, and at Jump8 (TMS_Jump8_), during the decision period. The figure also illustrates the three index-finger flexions required to provide a response. **(D) Time course of the sessions.** Participants completed, in each session, 4 training blocks (2 difficulty levels), the main Tokens task in 16 (TMS_Hand_) or 18 (TMS_Hand_) blocks, and 2 blocks of a simple reaction time (SRT) task. A brief discovery block was included only in the first (TMS_Hand_) session. **(E) Schematic representation of urgency modelling steps. Left panel:** The sum of log-likelihood ratios (SumLogLR) were computed within bins of increasing Tokens-task decision durations, separately for each precue × block-type condition. **Middle panel:** decision SumLogLR values were then converted into urgency using a regularised inverse. **Right panel:** urgency was modelled as a linear function of decision duration for each participant and condition, yielding 3 parameters: initial urgency (U-Intercept), urgency at Jump8 (U-Jump8; TMS time), and urgency growth rate (U-Slope).

#### Success probability and trial types

A key feature of the Tokens task is that it allows modulating the momentary distribution of tokens across the three circles over time, determining the “success probability” *p_i_(t).* The success probability is defined as the probability that a given choice (i.e., left or right lateral circle) would ultimately be correct at each time point *t* of trial *i*. This value depends on the momentary distribution of tokens across the three circles: the number of tokens already present in the left (N_L_) and right (N_R_) lateral circles, and the number remaining in the central circle (N_C_). If at any given moment, the left (L) circle contains N_L_ tokens, the right (R) one contains N_R_ tokens, and N_C_ tokens remain in the central (C) circle, then the probability that the left response is ultimately the correct one (i.e., the success probability of choosing left) can be computed as follows, assuming that “correct” means ending with a majority (≥ 8 tokens):

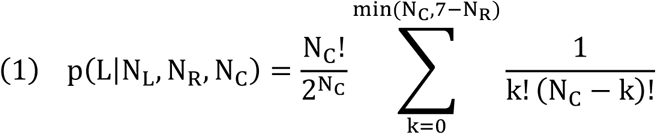

In words, this sums the probabilities of all outcomes where the right circle finishes with at most 7 tokens (hence left has the majority). Computing this probability for each of the 15 token jumps allowed us to characterize the temporal profile of success probability *p_i_(t)* for each trial. Although the individual token jumps appeared random to participants, the sequence of trials included distinct trial types, each defined by specific success probability dynamics (see Fig. 1A, upper part of middle panel). In the “obvious” trials, the *p_i_(t)* was above 0.7 after Jump_3_ and above 0.8 after Jump_5_, indicating that the initial token jumps consistently favored the correct choice. In the “misleading” trials, the *p_i_(t)* was below 0.4 before Jump_4_, reflecting an initial bias toward the incorrect option, which was subsequently reversed by later token jumps favoring the correct choice. For the “ambiguous” trials, we implemented four distinct subtypes in which the *p_i_(t)* remained between 0.3 and 0.66 until Jump_6_, Jump_8_, Jump_10_ or Jump_12_ (i.e., Ambiguous_6_, Ambiguous_8_, Ambiguous_10_, Ambiguous_12_ respectively), and above 0.6 after this jump. In these trials, the initial token jumps were balanced between the two lateral circles to keep the *p_i_(t)* close to chance level (i.e., 0.5). For example, in Ambiguous_6_ trials, the tokens jumped alternatively to the correct and incorrect lateral circles until Jump_6,_ such that the number of tokens was equal in both lateral circles after each even-numbered jump (i.e., after Jumps 2, 4, and 6), and differed by one after each odd-numbered jump (i.e., after Jumps 1, 3 and 5). Note that only Ambiguous_8_ trials were used for the first five participants; the full set of ambiguous subtypes was subsequently introduced for the remaining fifteen participants to increase variability in evidence dynamics. Finally, a subset of trials was entirely random, involving only trials with unpredictable success probability profiles *p_i_(t).* These trials were included to increase variability in the pattern of token jumps but were not considered in our analyses.

#### Experimental conditions

In the present study, we sought to dissociate control-driven and reward-driven urgency regulation and to examine how these modes are expressed in the motor system. To this end, we adapted the Tokens task with two crossed manipulations: one targeting control-driven regulation by varying the required level of control in separate blocks, and one targeting reward-driven regulation by varying the reward value of single trials.

To promote control-driven adjustments of urgency, participants performed the Tokens task in two block types that differed in their proportion of trial types (see Fig. 1A, right panel). In the “Low control” (Ctrl_Low_) blocks, there was a higher proportion of obvious trials (35%) and fewer misleading trials (15%), whereas in the “High control” (Ctrl_High_) blocks, there were fewer obvious trials (20%) and more misleading trials (30%). Both block types contained the same proportion of ambiguous (40%) and random (10%) trials. Although they were not informed about the existence of these specific trial types, participants were told in advance whether the upcoming block would require a low or high control level, and they were instructed to prioritize speed in Ctrl_Low_ blocks and accuracy in Ctrl_High_ blocks. Note however that to prevent participants from adopting overly cautious strategies and preserve a sufficient sense of decision urgency across block types, participants were informed that a bonus (+50 cents) could be obtained for sufficiently rapid and accurate decisions (see next section for details).

To promote reward-driven adjustments of urgency, we added a precue at the beginning of each trial, indicating the monetary reward participants could obtain for making a correct decision on that trial. Two types of precue were used (see Fig. 1A, lower part of middle panel): a “low reward” (Rew_Low_) precue (1 cent), shown as an image of a single coin, and a “high reward” (Rew_High_) precue (10 cents), shown as an image of a stack of coins. We expected higher rewards to increase the value of accuracy and thus bias regulation toward greater caution, resulting in lower urgency and later commitment when maximizing reward depends on avoiding errors. Crossing block type (Ctrl_Low_ vs. Ctrl_High_) with precue type (Rew_Low_ vs. Rew_High_) yielded four within-subject conditions (Ctrl_Low_–Rew_Low_, Ctrl_Low_–Rew_High_, Ctrl_High_–Rew_Low_, Ctrl_High_–Rew_High_), with the lowest urgency predicted in Ctrl_High_–Rew_High_ and the highest in Ctrl_Low_–Rew_Low_.

#### Sequence of events in a trial

Figure 1B illustrates the sequence of events within a single trial of the Tokens task. Each trial began with a blank screen of variable duration (250-650 ms) to signal trial onset. This was followed by the presentation of the reward precue displayed for 1000 ms and the pre-decision phase, during which the three circles appeared at the center of the screen, with the central circle containing 15 tokens. Subsequently, the tokens began to jump one by one from the central circle to one of the two lateral circles at a fixed rate of one jump every 200 ms. As described previously, participants were instructed to predict which lateral circle would ultimately receive the majority of tokens and to indicate their choice by performing three consecutive flexions of the left or right index finger. The first flexion triggered a Pac-Man animation, which advanced with each flexion by “eating” a dot, visually illustrating the motor sequence. Note that participants performed three rather than just one flexion for the purpose of another analysis addressed in a separate study. This aspect of the task is not critical for this study, which focused on TMS probes preceding the participants’ response. After these movements were made, the remaining tokens continued to jump until the central circle was empty. Each trial ended with performance feedback: the selected circle turned green for a correct choice and red for an incorrect one. In addition, the corresponding monetary reward was displayed below the central circle, together with a specific auditory cue: the reward was written in green for correct trials (i.e., “+1 cent” or “+10 cents”), whereas “+0 cent” was displayed in black for incorrect trials. The feedback screen lasted for 500 ms and then disappeared at the same time as the tokens and the 3 circles, denoting the end of the trial. This phase was followed by a blank screen that lasted during a variable intertrial interval (1800-2000 ms).

At the end of each block (40 trials), participants viewed a 3000 ms feedback screen displaying the total reward (in cents) for that block, corresponding to the sum of the reward earned on each trial plus a potential 50-cent bonus. This bonus depended on performance across the entire block: it was granted if the mean reaction time (RT) was below 2100 ms and accuracy exceeded 55% for each reward precue type. The block-level feedback also included a brief message indicating whether the bonus had been obtained and summarizing performance (e.g., “Congrats, you got the bonus!”, “Good speed, but bad accuracy”, or “Be faster”). This allowed participants to evaluate and adjust their decision strategy for the next block.

### 2.3 Transcranial Magnetic Stimulation

Participants received TMS in two separate sessions on different days to elicit MEPs either in hand muscles (TMS_Hand_ session) or in leg muscles (TMS_Leg_ session. see Fig. 1C, right panel).

In both sessions, we first identified the optimal coil position (i.e., hotspot) on primary motor cortex (M1) to elicit MEPs in the reference muscle, namely, the First Dorsal Interosseus (FDI) and the Tibialis Anterior (TA) in the TMS_Hand_ and TMS_Leg_ sessions, respectively. These hotspots were carefully marked on a head cap placed on the participant’s scalp to ensure consistent coil positioning throughout the experimental session (Yadav et al., 2025). We then estimated the individual resting motor threshold (rMT), defined as the minimal intensity required to evoke MEPs of 50 μV peak to peak on 5 out of 10 consecutive trials in the contralateral muscle of reference (FDI or TA) (Rossini et al., 2015). In each session, the stimulation intensity was always set at 115% of the individual rMT and maintained at this level for the entire experiment. One participant was excluded because he was unable to tolerate the stimulation intensity required during the rMT procedure. Moreover, 1 participant was tested but not included in the analyses because MEPs progressively vanished during the session, precluding consistent measurements. Finally, 3 more participants were excluded because of technical issues in which TMS pulses were either not delivered or delivered at incorrect time points. Hence, a total of 20 participants were included in the final sample.

In both sessions, TMS pulses were applied at two different times during task blocks (see Fig. 1C, left panel). First, baseline MEPs were elicited within blocks (TMS_Baseline-in_) during the trial onset, with pulses occurring 250–650 ms before the reward precue. These baseline measures provide a probe of corticospinal excitability when participants were still at rest but already engaged in the task context. Second, and most importantly, TMS pulses were applied during the deliberation process, specifically at the 8^th^ token jump (TMS_Jump8_), corresponding to 1400 ms after the first token jump, allowing us to assess changes in corticospinal excitability during decision making, as a function of changes in urgency. Finally, some pulses were delivered at three time points outside the task blocks (TMS_Baseline-out_) to obtain baseline measures of corticospinal excitability when subjects were completely at rest with the eyes open (Fig. 1D).

TMS was applied across all trial types, ensuring that participants could not associate a particular trial type with TMS occurrence. However, MEPs recorded during obvious, misleading and random trials were excluded from the main analyses, as these trials typically elicited responses before Jump_8_. Our primary focus was therefore on ambiguous trials, in which the token distribution remained balanced between the two lateral circles until Jump_8_. This ensured that participants’ responses generally occurred after Jump_8_ and minimized any strong accumulation of sensory evidence before TMS delivery. Consequently, observed changes in motor excitability at this timing and differences between experimental conditions could not be attributed to sensory evidence differences. More details specific to each session are provided below.

#### TMS_Hand_ session

In this session, TMS was applied using a double-coil protocol, wherein both M1 FDI hotspots were stimulated with a 1 ms inter-pulse interval (see Fig. 1C, right panel) (Grandjean et al., 2018; Vassiliadis et al., 2020). This approach allows to elicit near-simultaneous MEPs in finger muscles of both hands and has been shown to produce MEPs comparable in amplitude and reliability to those obtained with classic single-coil protocols, while requiring only half the time, as MEPs are acquired from both hands within a single trial (Grandjean et al., 2018; Vassiliadis et al., 2018; Quoilin et al., 2019; Derosiere et al., 2020).

Stimulations were delivered via two small figure-of-eight coils (wing internal diameter of 35 mm), each connected to a monophasic stimulator (one Magstim 200 and one Magstim Bistim^2^; Magstim, Whitland, Dyffed, United Kingdom). The coils were positioned tangentially over the left and right side of the scalp, with handles pointing backward and oriented laterally at a 45° angle away from the midline, inducing a postero-anterior current in the cortex. On average, the rMT for left and right M1 FDI hotspots was 43 ± 8.7% and 43 ± 9.3% of maximum stimulator output (MSO), respectively. Stimulation applied over the hotspot for the FDI (task agonist) also elicited reliable MEPs in the Abductor Pollicis Brevis (APB), a nearby but task-irrelevant muscle.

#### TMS_Leg_ session

During this session, TMS was applied over the TA hotspot of the right M1 (see Fig. 1C, right panel), using a batwing coil (D-B80 Magpro coil) connected to a biphasic stimulator (Magpro X100 Stimulator; Magventure, Farum, Denmark), following the protocol described in Derosiere et al., 2022. This specific coil was selected for its ability to effectively stimulate deep cortical structures within the interhemispheric fissure, where lower limb motor representations are located. These deep regions are not easily accessible with conventional figure-of-eight coils, which primarily activate superficial cortical layers (Deng et al., 2014). To further optimize stimulation efficacy, biphasic pulses were used as they have been shown to better recruit deep neural populations compared to monophasic pulses (Niehaus et al., 2000). The coil was first positioned tangentially on the scalp vertex, with the handle pointing backward and aligned with the midline, to induce a current with a posteroanterior direction in the cortex. It was then gradually rotated counterclockwise to direct the magnetic field toward the right M1 leg representation, eliciting reliable MEPs in the left TA. As for the TMS_Hand_ session, the coil position was carefully marked on a head cap placed on the participant’s scalp, providing a reference point throughout the session. The rMTs at right M1 TA hotspot was 66 ± 13.4% of MSO. In addition to eliciting MEPs in the left TA by stimulating its hotspot, TMS also evoked reliable MEPs in other muscles of the left leg, including the Vastus Lateralis (VL) and Rectus Femoris (RF) muscles. Because the Tokens task required left or right index finger movements, these left leg MEPs could either occur on the chosen side (during left-hand choices) or the unchosen side (during right-hand choices).

#### Electromyography (EMG) recording

EMG activity was recorded using disposable surface electrodes (BlueSensor NF-50-K, Ambu, Ballerup, Denmark) positioned over the target muscles in the TMS_Hand_ session (left and right FDI and ABP) and TMS_Leg_ session (left TA, VL, and RF). A ground electrode was also placed over the left ulnar styloid process in both sessions. For each trial, EMG signals were recorded for 3200 ms, beginning 200 ms before the TMS pulse. Raw EMG signals were amplified (gain of 1000), band-pass filtered (10 to 500 Hz) and notch filtered (50 Hz) online (NeuroLog, Digitimer, UK), and digitized at 2000 Hz for offline analysis. Experimenters continuously monitored the signals throughout the session and instructed participants to relax whenever background muscle activity was detected.

### 2.4 Time course of the sessions

As mentioned above, the participants completed two sessions in the laboratory, with the structure of each one illustrated in Figure 1D. Participants always completed the TMS_Hand_ session first, followed the next day or, at most, within one week by the TMS_Leg_ session. Both sessions were conducted at the same time of day to control for circadian variations in corticospinal excitability and the session order was kept constant across participants, as no direct comparisons between the two sessions were planned. The structure of the two sessions was identical except for a short discovery block in the TMS_Hand_ session. This block comprised 10 trials without reward precues and included a balanced distribution of trial types, allowing participants to familiarize themselves with the fundamental principles of the Tokens task before the main experiment began. In both sessions, participants then completed four training blocks of 15 trials each, in which the reward precue (Rew_Low_, Rew_High_) was implemented. Each session included two training blocks per control level (Ctrl_Low_, Ctrl_High_), presented in an alternating order, with the first block type counterbalanced across participants. TMS was introduced only during the second round of training blocks, with single pulses delivered either over the FDI hotspot, in the TMS_Hand_ session, or the TA hotspot, in the TMS_Leg_ session. This step allowed participants to become familiar with the sensation of TMS during task execution. Once training was completed, the main experimental phase began. This phase consisted of 16 blocks in the TMS_Hand_ session and 18 blocks in the TMS_Leg_ session. The number of blocks was greater in the TMS_Leg_ session because only one leg (one M1) was stimulated at a time, which required additional blocks to obtain enough MEPs on both the chosen and unchosen sides. Each session included an equal number of Ctrl_Low_ and Ctrl_High_ blocks, each consisting of 40 trials. Block types alternated, and the starting block was randomized across participants. Within each block, 80% of the trials (32 out of 40) were stimulated, while the remaining 20% (8 out of 40) did not involve stimulation to prevent anticipation of TMS pulses. Among the 32 stimulated trials, 9 MEPs were elicited at TMS_Baseline-in_, and 23 MEPs were elicited at TMS_Jump8_, including 14 ambiguous trials retained for analysis. Overall, this protocol yielded a total of 256 MEPs in the TMS_Hand_ session (72 MEPs at TMS_Baseline-in_ and 184 MEPs at TMS_Jump8_) and 252 MEPs in the TMS_Leg_ session (81 MEPs at TMS_Baseline-in_ and 207 MEPs at TMS_Jump8_). Each block lasted approximately 6-7 minutes, and a short break was provided between blocks to allow participants to rest. Finally, as mentioned above, MEPs were also elicited at TMS_Baseline-out_, that is, in between task blocks. Specifically, 20 MEPs were recorded at three distinct time points: before the first experimental block, halfway through the session, and after the last experimental block.

Each session ended with 2 blocks of a simple reaction time (SRT) task (20 trials each), which used the same visual display as the Tokens task described above. However, here, after briefly appearing in the central circle (500 to 900 ms), the 15 tokens jumped altogether simultaneously into one of the two lateral circles. A variable delay was introduced before the token jump to prevent participants from anticipating the jump. Crucially, in one block, the tokens always jumped to the left circle, and in the other, they always jumped to the right circle. The order of these two blocks was randomized across participants and sessions. Because the target circle remained constant within each block and was known in advance, participants simply had to respond as quickly as possible (without anticipating) by flexing the index finger corresponding to the target circle. This task involved no decision-making and served solely to estimate each participant’s median SRT for left and right index finger movements(Cisek et al., 2009; Derosiere et al., 2022). The total duration of a session was approximately 4 hours.

### 2.5 Data preprocessing and endpoint measures

#### Behavior in the Tokens task: urgency and performance-related endpoint measures

Behavioral data were acquired and analyzed using custom Matlab scripts (MathWorks, Natick, Massachusetts, USA).

##### Urgency-related endpoint measures

We exploited the urgency-gating framework to extract an urgency function for each reward precue × block type condition in each participant. To do so, we followed the procedure used in previous studies from our group (Derosiere et al., 2021, 2022). In brief, the task provides an estimate of the sensory evidence available at the time of commitment, that is, the amount of information on which the participant’s choice is based. This evidence was approximated after each token jump by a first-order estimate of the true success probability, known as the sum of log-likelihood ratios (SumLogLR):

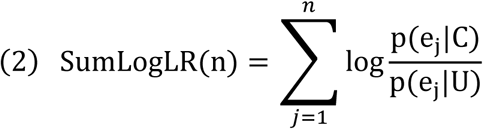

In this equation, *p(ej|C)* denotes the likelihood of observing token event *ej* (i.e., a token jumping either toward the chosen or the unchosen side) on trials in which the chosen side *C* is correct, and *p(ej|U)* is the likelihood of the same event *ej* on trials in which the unchosen side *U* is correct. Hence, the SumLogLR is proportional to the difference in the number of tokens favoring the chosen versus the unchosen lateral circle and therefore quantifies the strength of evidence at any given time.

For each participant and each reward precue × block type condition, we sorted trials into 10 quantiles of decision durations and computed, for each bin, the mean SumLogLR-at-decision (Fig. 1E, left panel). Because the proportion of trial types differed between Ctrl_Low_ and Ctrl_High_ blocks, we applied within-subject weighting to equalize the relative contribution of each trial type to the bin averages across conditions. Under the urgency-gating model, choices occur when the product of sensory evidence and urgency reaches a fixed threshold; therefore, the urgency at commitment is inversely related to the available evidence. The urgency was computed as follows: under a fixed threshold, we converted the (weighted) bin–wise SumLogLR–at–decision values to urgency using a regularized inverse (Fig. 1E, middle panel):

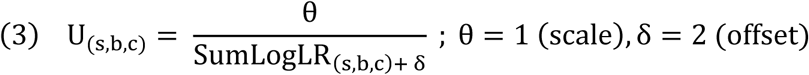

In this equation, *s* denotes the participant, *b* the decision duration bin and *c* the condition. ϴ fixes the scale of the urgency signal (units are arbitrary but constant across subjects/conditions), and 𝛿 is a small positive offset to prevent singularities when evidence is near zero and to keep urgency values positive and numerically stable. We then fitted, for each participant and each reward precue × block type condition, a linear function of urgency against decision duration. From this function, we extracted three parameters (Fig. 1E, right panel): the urgency intercept (*U-Intercept*; initial urgency), its predicted value at Jump_8_ *(U-Jump_8_)*, corresponding to the time of the TMS probes, and the slope (temporal growth) of urgency (*U-Slope)*. These three parameters (U-Intercept, U-Jump_8_ and U-Slope) served as our urgency-related endpoint measures.

##### Performance-related endpoint measures

Two main endpoint measures were used to assess how control-driven and reward-driven urgency regulation translated into Tokens-task performance. First, we considered the time participants took to commit to a choice, which we refer to as the *decision duration*. This measure corresponds to the interval between the first token jump (Jump_1_) and the onset of the first index finger flexion, minus the median SRT from the last two SRT blocks of the session. This yields an estimate of decision duration that is largely free of non-decisional sensorimotor delays (Cisek et al., 2009; Derosiere et al., 2019; Thura, 2020; Derosiere et al., 2022; Carsten et al., 2023). Second, we considered the *decision accuracy*, defined as the percentage of trials in which the participant selected the correct lateral circle. These two parameters (decision duration and decision accuracy) served as our performance-related endpoint measures. Note that all behavioral measures were computed based on data pooled across TMS conditions, blocks and sessions, as justified in the Supplementary Materials (see Suppl. Fig. 1A). Importantly, the first five participants who performed only Ambiguous_8_ trials were included, as performance-related results remained unchanged whether these participants were included or excluded from the analyses.

#### TMS probes in the Tokens task: broad modulation and surround inhibition

TMS-related data were acquired via EMG using Signal (Cambridge Electronic Design, Cambridge, UK) and extracted with a custom-written Signal script. The resulting EMG data were then analyzed using Matlab (MathWorks, Natick, Massachusetts, USA).

As described above, only MEPs elicited in ambiguous trials were included in the analysis. MEPs were obtained in two separate sessions, with hand muscles (FDI, APB) targeted in the TMS_Hand_ session and leg muscles (TA, VL, RF) targeted in the TMS_Leg_ session. For all these muscles, we extracted the peak-to-peak amplitude of MEPs elicited at each time of interest, namely at TMS_Baseline-out_, TMS_Baseline-in_ and TMS_Jump8_. MEPs from all trials were considered, independent of whether the correct or incorrect lateral circle was selected. At each time point, we discarded trials with excessive background EMG activity in the 200 ms pre-TMS window (Root Mean Square > 2.5 SD) and trials with extreme MEP amplitudes (>2.5 SD). Note that a trial was retained only if all recorded muscles passed these criteria (FDI and APB bilaterally for the TMS_Hand_ session; TA, VL, RF for the TMS_Leg_ session). Following preprocessing, an average of 199 trials (minimum of 33 trials per condition) and 230 trials (minimum of 29 trials per condition) remained at TMS_Jump8_ for the TMS_Hand_ and TMS_Leg_ sessions, respectively. Importantly, no participant was excluded from the TMS_Hand_ session after this procedure. However, in the TMS_Leg_ session, two participants had to be excluded: one due to insufficient residual trials after preprocessing (i.e., poor quality of EMG data) and another because the session could not be completed, resulting in insufficient number of trials for reliable analysis. Consequently, all analyses for the TMS_Leg_ session were conducted on the remaining 18 participants. Then, further processing was applied specifically to MEPs elicited at TMS_Jump8_ (i.e., 1400 ms after the first token jump). First, to ensure that MEPs reflected deliberation rather than movement execution, we retained only trials in which the first index finger flexion occurred at least 200 ms after the TMS pulse (i.e., after 1600 ms). Then, to eliminate variability in MEP amplitude that could arise from differences in decision duration across trials or conditions, we applied a strict RT-matching procedure for trial selection based on a single automatically selected window (Murphy et al., 2016; see Supplementary Materials and Suppl. Fig. 1B for more details on the procedure). For the TMS_Hand_ session, the selected window [1900-2400ms] allowed exactly 12 trials per subject × condition). For the TMS_Leg_ session, the selected window [1700–2400ms] also included 12 trials per subject × condition, and an additional constraint enforced a 50:50 split of left vs right choices (≥6 per side) to prevent bias from unilateral recording. Within each window, trials were chosen deterministically, prioritizing those closest to the window center. This approach ensured that TMS_Jump8_ MEPs were compared within a matched and standardized RT range across all conditions, thereby eliminating timing-related confounds while maintaining balanced datasets.

Remaining MEPs at TMS_Jump8_ were expressed in percentage of the mean MEP amplitude at TMS_Baseline-in_. These normalized MEPs at TMS_Jump8_ (%MEP_J8_) were used to compute the probes of broad modulation and surround inhibition. For broad modulation, we considered the %MEP_J8_ from the TMS_Leg_ session averaged across the three leg muscles (TA, VL, RF) on each trial, irrespective of the chosen circle, with this %MEP_J8-Leg_ serving as a probe of wide-ranging corticospinal modulation. We also assessed whether broad modulation could be observed for the prime-mover in the task by considering MEPs in the FDI (%MEP_J8-FDI_) as in theory, broad modulation should affect corticospinal excitability in all muscles, not only in the most distant ones. Then, for surround inhibition, we used the %MEP_J8_ recorded from the APB and FDI muscles (%MEP_J8-APB_ and %MEP_J8-FDI_, respectively), where we expected a specific suppression of %MEP_J8-APB_ compared to %MEP_J8-FDI_, on the response side (chosen circle). This effect is taken as evidence of surround inhibition affecting task-irrelevant muscles (i.e., the APB) that lie close to (surround) the prime mover (i.e., the FDI). To quantify this specific suppression of APB, we computed a delta (Δ) index on each trial, reflecting the difference between %MEP_J8-APB_ and %MEP_J8-FDI_ on the specific trial (APB − FDI) (Leodori et al., 2019; Thirugnanasambandam et al., 2020). Under surround inhibition, this Δ% MEP_J8-APB_ should be negative, with more negative values indicating stronger surround inhibition.

As a reminder, our main hypothesis was that control-driven urgency regulation (block type effect) would primarily manifest as changes in broad modulation of motor excitability, whereas reward-driven urgency regulation (precue type effect) would primarily manifest as changes in surround inhibition.

### 2.6 Statistical analyses

Statistical analyses were conducted using Jamovi 2.5.3 (The jamovi project, Sydney, Australia). Urgency-related measures (U-Intercept, U-Jump_8_ and U-Slope), performance-related measures (decision duration and decision accuracy) and TMS probes of broad modulation (%MEP_J8-Leg_ and %MEP_J8-FDI_) and surround inhibition (Δ%MEP_J8-APB_) were analyzed at the single-trial level using linear and generalized (decision accuracy) mixed-effects models.

All measures were analyzed with *precue type* (Rew_Low_/Rew_High_) and *block type* (Ctrl_Low_/ Ctrl_High_) as fixed factors, and *UPPS_Urgency_* as a covariate to assess the relationship between endpoint measures and urgency dimension of impulsivity. Participant was included as a random effect (i.e., random intercept). For the performance-related behavioral measures, *trial type* (obvious, misleading, ambiguous) was also added as a fixed factor. Note that obvious trials were not entered in the analysis of decision accuracy due to near-ceiling performance causing model overfitting. Moreover, as explained before, the TMS probes were only considered for ambiguous trials. Post-hoc comparisons following significant interactions were adjusted using Tukey corrections. When interactions involved UPPS_Urgency_, we further examined simple effects by estimating model predictions at three representative levels of the covariate: “low” urgency (mean UPPSUrgency minus 1 SD), “average” urgency (mean), and “high” urgency (mean plus 1 SD)(Bauer & Curran, 2005). Finally, further post-hoc associations were examined using Student’s t-tests, with p-values corrected for multiple comparisons when appropriate (p-corrected). Statistical significance was set at p < 0.05, and all descriptive data are reported as estimated marginal means ± standard errors (SE) unless otherwise specified.

## 3. Results

### Behavior in the Tokens task: urgency and performance-related endpoint measures

#### Urgency-related endpoint measures

We first examined behavior of participants using the urgency–gating framework. Urgency functions are shown on Figure 2A with all subjects pooled together on the left-most panel (n=20); for visualization purpose, we also show the urgency functions for three representative participants displaying low (left-middle panel), average (right-middle panel) or high (right-most panel) *UPPS_Urgency_* scores. Urgency parameters (i.e., U-Intercept, U-Jump_8_ and U-Slope) were extracted for each experimental condition in each participant, following the procedure described in the method section. This yielded four measures (2 *block type* x 2 *precue type*) for each urgency parameter in each participant. Here, we focus on U-Jump8, which captures the urgency level at the timing of the TMS probes and integrates both baseline urgency and its temporal growth (see insets of Fig. 2A); analyses of U-Intercept and U-Slope led to similar observations and are presented in the Supplementary Materials (see Suppl. Fig. 2)

**Figure 2.**
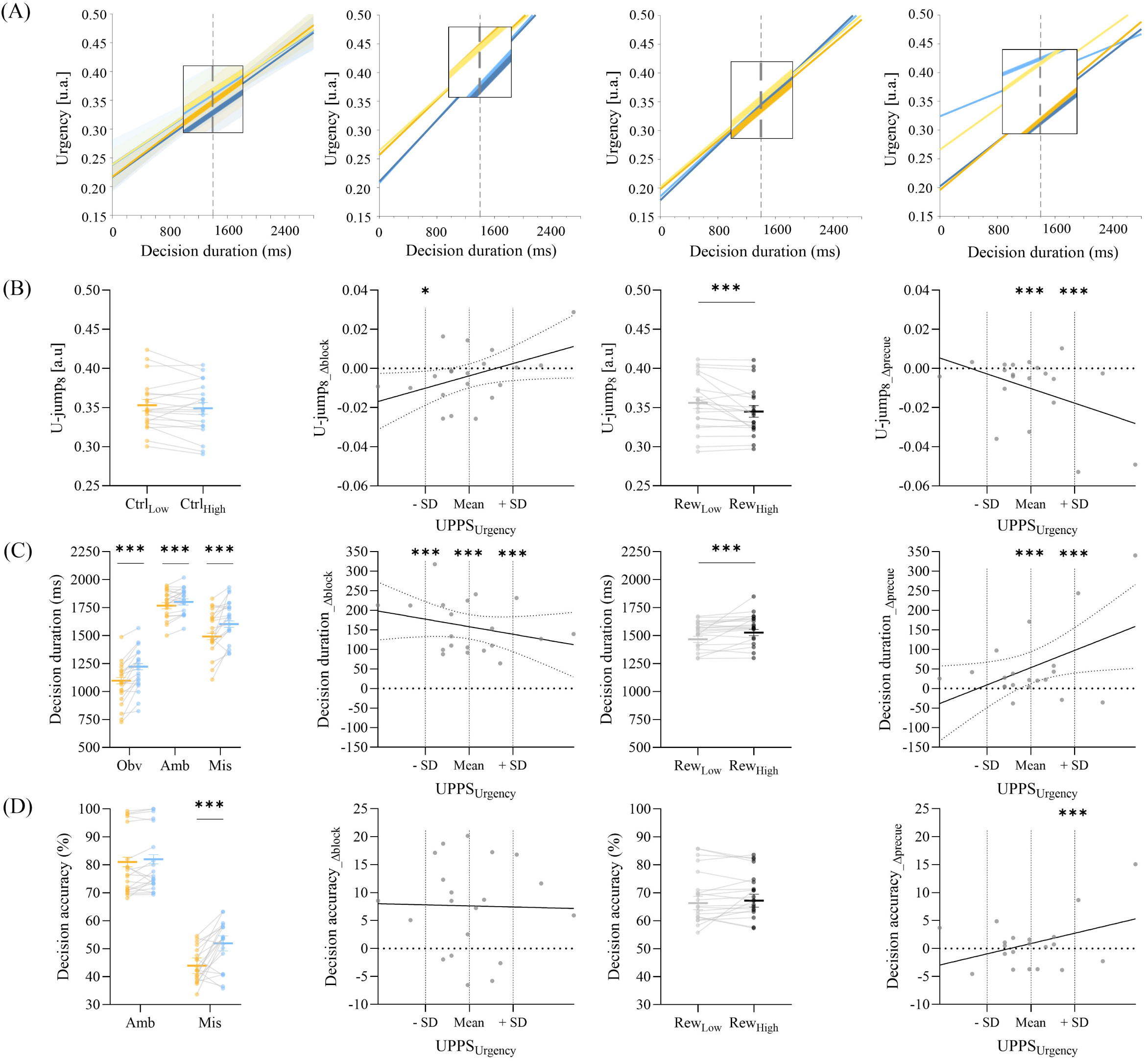
Main results on behavioral data. **(A) Urgency functions as a function of decision duration.** Urgency functions are presented in Ctrl_Low_ (yellow) and Ctrl_High_ (blue) blocks involving Rew_Low_ and Rew_High_ precues (light and dark colors, respectively). **Left-most panel**: Urgency functions are presented for all participants together. Shaded bands indicate fit uncertainty. **Middle three panel:** The same fits are presented for three representative participants displaying low (Left-middle panel), average (Right-middle panel) and high (Right-most panel) UPPS_Urgency_ scores. **(B) U-Jump8, (C) Decision duration and (D) Decision accuracy results**. **Left-most panel:** results are shown as a function of *block type* with Ctrl_Low_ blocks in yellow and Ctrl_High_ blocks in blue. **Left-middle panel:** All behavioral measures are presented as a delta (Δblock: Ctrl_High_ – Ctrl_Low_) for low, average and high representative UPPS_Urgency_ values. **Right-middle panel:** Behavioral parameters are presented as a function of precue type with Rew_Low_ in light and Rew_High_ in dark**. Right-most panel:** All behavioral measures are presented as a delta (Δprecue: Rew_High_ – Rew_Low_) for low, average and high representative UPPS_Urgency_ values. *Bold horizontal lines represent estimated marginal means, error bars indicate standard error, gray lines connect within–participant values, and dots represent individual participants.*: p-value < 0.05; ***: p-value < 0.001*

The linear mixed-effects model showed no evidence for an overall difference in U-Jump8 between Ctrl_Low_ and Ctrl_High_ blocks (F_(1, 54)_ = 1.66, p = 0.203; Fig. 2B, left-most panel). However, there was a *block type × UPPS_Urgency_* interaction (F_(1, 54)_ = 4.39, p = 0.041), indicating that block-related adjustments depended on the impulsivity score at the UPPS_Urgency_ subscale. To probe this interaction, we ran a post-hoc analysis examining the simple effects of *block type* on U-Jump8 (illustrated as a delta [Δblock: Ctrl_High_ minus Ctrl_Low_] in Fig. 2B, left-middle panel) at three representative UPPS_Urgency_ values, with “low” corresponding to the mean UPPS_Urgency_ minus 1 SD, “average” corresponding to the mean UPPS_Urgency_, and “high” corresponding to the mean UPPS_Urgency_ plus 1 SD. At low UPPS_Urgency_ values, U–Jump8 was lesser in Ctrl_High_ than Ctrl_Low_ blocks (F(1, 54) = 5.73, p = 0.020), consistent with a down-regulation of urgency in Ctrl_High_ blocks among less impulsive participants. This slowing was not observed at average or high UPPS_Urgency_ values where the *block type* effect was not significant (all F < 1.66, all p > 0.203). This heterogeneity likely accounts for the absence of a *block type* effect at the group level. In contrast, there was a main effect of *precue type* (F_(1, 54)_ = 12.24, p < 0.001; Fig. 2B, right-middle panel), with U–Jump8 being overall lower following Rew_High_ precue than Rew_Low_ precue. Interestingly, this effect also depended on impulsivity, as indicated by a *precue type × UPPS_Urgency_* interaction (F_(1, 54)_ = 6.21, p = 0.016). Post-hoc simple–effect tests of *precue type* (illustrated as a delta [Δprecue: Rew_High_ minus Rew_Low_] in Fig. 2B, right-most panel) at representative UPPS_Urgency_ values revealed a significant precue effect at average and high UPPS_Urgency_ scores (both F_(1, 54)_ > 12.24, both p < 0.001) but not for individuals displaying the lowest scores (F_(1, 54)_ = 0.489, p = 0.487).

Taken together, our results on U-Jump8 indicate distinct adjustment patterns as a function of participants’ impulsivity scores (see Supplementary Materials and Suppl. Fig. 2A for comparable results on U-Intercept). Control-driven down-regulation of urgency was primarily observed among less impulsive participants, suggesting that only this subgroup effectively adjusted urgency to increased task demands in Ctrl_High_ blocks. By contrast, reward-driven down-regulation of urgency following Rew_High_ precues emerged at the group level but was weaker among less impulsive participants, implying that the reward effect on urgency was largely driven by the more impulsive individuals.

#### Performance-related endpoint measures

We then examined how the urgency adjustments described above translated into modulations of decision duration and accuracy in the Tokens task. Here, we focus on results that inform the performance consequences of these urgency adjustments, including effects involving UPPS_Urgency_; all other results, which largely confirmed the expected effectiveness of our manipulations, are reported in the Supplementary Materials.

Decision durations showed the expected slowing in Ctrl_High_ relative to Ctrl_Low_ blocks (main effect of *block type*: F_(1, 23083.1)_ = 290.32, p < 0.001). This slowing was present for each trial type (*block type x trial type* interaction: F_(2, 23083)_ = 35.12, p < 0.001; Fig. 2C, left-most panel; see also Supplementary Materials for more details on results involving *trial type*) but varied with UPPS_Urgency_ (*block type x UPPS_Urgency_* interaction: F_(1, 23083.1)_ = 5.66, p = 0.017). Post-hoc simple-effects tests indicated significant slowing at low, average, and high UPPS_Urgency_ (all χ² > 107, all p < 0.001; all positive Δblock values in Fig. 2C, left-middle panel), but the magnitude of this effect decreased as UPPS_Urgency_ increased. Turning to the effect of the *precue type*, decision durations were slower following Rew_High_ than Rew_Low_ precues (main effect of *precue type*: F_(1, 23083)_ = 127.28, p < 0.001; Fig. 2C, right-middle panel), but again this effect depended on UPPS_Urgency_ (*precue type x UPPS_Urgency_* interaction: F_(1, 23083)_ = 114.06, p < 0.001). Post-hoc simple-effects analyses of *precue type* (illustrated as Δprecue in Fig. 2C, right-most panel) at representative UPPS_Urgency_ values showed a significant slowing following Rew_High_ precue at average and high UPPS_Urgency_ scores (both χ^2^(1) > 127.26, both p < 0.001), but not at low UPPS_Urgency_ (χ²(1) = 0.178, p = 0.673).

Continuing with decision accuracy, there was a *block type × trial type* interaction (χ²_(1)_ = 11.17, p < 0.001; Fig. 2D, left-most panel), indicating a significant enhancement in the Ctrl_High_ blocks that was only present in the most challenging (misleading) trials (p < 0.001), but this effect did not interact with *UPPS_Urgency_* as the triple interaction was not significant (χ²(1) = 0.08, p = 0.783; see the consistent Δblock across *UPPS_Urgency_* on Fig. 2D, left-middle panel). In contrast, *UPPS_Urgency_* influenced the way participants adjusted their accuracy according to the *precue type*. As such, there was no main effect (χ²_(1)_ = 1.08, p = 0.299; Fig. 2D, right-middle panel) but a significant *precue type x UPPS_Urgency_* interaction (χ²_(1)_ = 6.53, p = 0.011). Simple–effect tests of *precue type* (Δprecue on Fig. 2D, right-most panel) at representative UPPS_Urgency_ values revealed a significant *precue type* effect at high UPPS_Urgency_ scores (χ^2^(1) =6.71, p = 0.010), but not at average and low impulsivity values (both χ^2^(1) < 1.08, both p > 0.299).

Hence, control-driven decision slowing was observed in all participants, but its magnitude decreased as UPPS_Urgency_ increased, consistent with the U-Jump8 findings. However, this control-driven slowing did not translate into a selective accuracy benefit in Ctrl_High_ blocks among less impulsive participants, despite their greater slowing. In line with the U-Jump8 results, reward-driven decision slowing was weaker in less impulsive participants and was accompanied by a more limited gain in decision accuracy than in more impulsive individuals.

#### TMS probes in the Tokens task: Broad modulation and Surround inhibition

The behavioral patterns described in the previous section confirm that, as intended, urgency and the resulting performance in the Tokens task varied both with control demands and reward context. Moreover, we observed dissociable adjustment profiles across impulsivity levels: less impulsive participants showed stronger control-driven adjustments, whereas more impulsive participants exhibited more pronounced reward-driven adjustments of urgency. In this section, we examined whether these behavioral adjustments were accompanied by specific changes in corticospinal excitability, and whether these changes were also related to impulsivity. To this end, we focused on the TMS probes at Jump8 defined in the Methods: %MEP_J8-Leg_ and %MEP_J8-FDI_ as probes of broad modulation and Δ%MEP_J8-APB_ as a probe of surround inhibition.

##### Broad modulation

Broad modulation was primarily evaluated based on %MEP_J8-Leg_, following the rationale that any global change in motor excitability should be detectable in task-irrelevant muscles located far from the task agonist, such as leg muscles during a task involving finger muscles. As explained in the Materials and Methods section, all subsequent analyses based on %MEP_J8-Leg_ were performed on 18 participants. One sample t-tests confirmed the presence of a significant broad modulation, manifesting as a broad facilitation, of %MEP_J8-Leg_ with respect to baseline (100%) in all conditions of the Tokens task (all t > 2.89, all p-corrected < 0.02; Fig. 3A, left panel). The linear mixed-effects analysis did not reveal any main effect, whether considering *block type* (F_(1, 2568)_ = 1.49, p = 0.222) or *precue type* (F_(1, 2568)_ = 0.934, p = 0.334). The absence of a precue type effect was confirmed by a Bayesian ANOVA, which provided evidence in favor of the null model (BF_10_ = 0.25), indicating comparable levels of broad facilitation across precue types. Interestingly, and similarly to the effects found on urgency and decision behavior, we observed a significant *block type × UPPS_Urgency_* interaction on %MEP_J8-Leg_ (F_(1, 2568)_ = 6.99, p = 0.008). Post-hoc simple–effect tests of *block type* (illustrated as Δblock in Fig. 3A, middle panel) at representative UPPS_Urgency_ values revealed significantly lower facilitation of %MEP_J8-Leg_ in Ctrl_High_ than Ctrl_Low_ blocks in participants displaying low UPPS_Urgency_ values (F_(1, 2568)_ = 7.47, p = 0.006), but not at average (F_(1, 2568)_ = 1.49, p = 0.222) and high UPPS_Urgency_ values (F_(1, 2568)_ = 1.01, p = 0.314). Hence, less impulsive individuals, who were also the only ones showing a control-driven down-regulation of U-Jump8 in Ctrl_High_ blocks were also those showing an attenuation of broad facilitation. We specifically examined this association between control-driven changes in broad modulation (%MEP_J8-Leg_Δblock_) and control-driven changes in urgency at the time of the TMS probe (U-Jump_8_Δblock_). Interestingly, we observed a positive correlation: participants who showed a larger reduction in broad facilitation in Ctrl_High_ blocks also exhibited a stronger down-regulation of U-Jump_8_ in these blocks compared with Ctrl_Low_ blocks (Pearson’s r = 0.407, p = 0.047; Fig. 3A, right panel). Taken together, these findings suggest that broad facilitation constitutes a motor implementation of control-driven urgency regulation, whose engagement is progressively reduced with increasing impulsivity.

**Figure 3.**
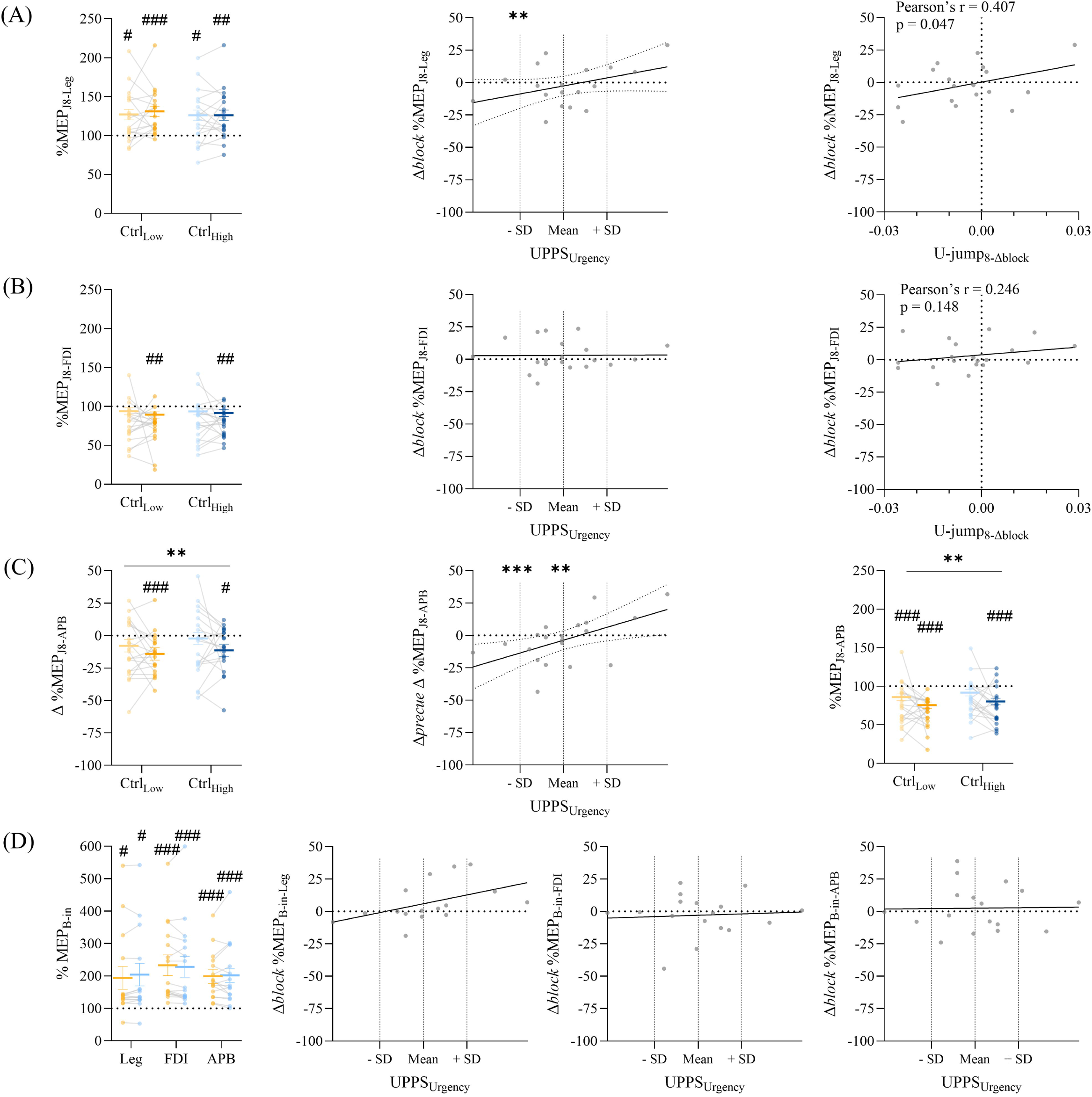
Main results on TMS probes. All results are presented as a function of block type with Ctrl_Low_ block in yellow and Ctrl_High_ block in blue, for the two precue levels with Rew_Low_ in light and Rew_High_ in dark. **(A) %MEP_J8-Leg_ and (B) %MEP_J8-FDI_ on chosen side. Left panel:** Results are expressed in percentage of baseline, with one-sample t-test comparisons to 100% indicated by # symbols. **Middle panel:** MEPs are presented as a delta (Δblock: Ctrl_High_ – Ctrl_Low_) for low, average and high representative UPPS_Urgency_ values. **Right panel:** MEPs are presented as a delta (Δblock = Ctrl_High_ – Ctrl_Low_) as a function of corresponding change in block type for urgency at Jump8 (Δblock = Ctrl_High_ – Ctrl_Low_)**. (C) Δ%MEP_J8-APB_ on chosen side. Left panel**: Results for Δ%MEP_J8-APB_ (APB – FDI) are presented with comparison one-sample t-test comparison to 0, with negative values indicating smaller MEPs in APB than FDI, indicated by # symbols. **Middle panel:** Δ%MEP_J8-APB_ (APB – FDI) is presented as a delta (Δprecue: Rew_High_ – Rew_Low_) for low, average and high representative UPPS_Urgency_ values. **Right panel:** %MEP_J8-APB_ (without FDI) are expressed in percentage of baseline, with one-sample t-test comparison to 100% indicated by # symbols. **(D) %MEP_B-in._ Left panel:** %MEP_B-in_ for each probe (Leg, FDI and APB) is expressed in percentage of %MEP_B-out_, with one-sample t-test comparison to 100% indicated by # symbols, for the two block types, Ctrl_Low_ and Ctrl_High_. **Three middle panel:** Baseline-in MEPs are presented as a delta (Δblock: Ctrl_High_ – Ctrl_Low_) for low, average and high representative UPPS_Urgency_ values, for each TMS probe separately (Leg at the left, FDI in the middle and APB at the right). *Bold horizontal lines represent estimated marginal means, error bars indicate standard error, gray lines connect within–participant values, and dots represent individual participants. **: p-value < 0.01; ***: p-value < 0.001 for mixed models. #: p-value < 0.05; ##: p-value < 0.01; ###: p-value < 0.001 for Student’s t-test*.

In theory, a broad modulation account should impact corticospinal excitability across muscles, not only in the most distant ones. Hence, it would be reasonable to also expect an effect on %MEP_J8-FDI_ in the prime mover, at least in less impulsive participants as observed for %MEP _J8-Leg_. However, when considering %MEP_J8-FDI_ on the chosen side, we observed no main effect of *block type* (F_(1, 934)_ = 0.15, p = 0.703) or *precue type* (F_(1, 934)_ = 1.22, p = 0.269; Fig. 3B, left panel). We also found no *block type × UPPS_Urgency_* interaction (F_(1, 934)_ = 1.45, p = 0.228). Together, these results indicate no decrease in FDI excitability in Ctrl_High_ relative to Ctrl_Low_ blocks across *UPPS_Urgency_* values, as reflected in Fig. 3B (middle panel), with uniformly null %MEP_J8-FDI_Δblock_ values. There was also no relationship between %MEP_J8-FDI_Δblock_ and block-related U-Jump_8_Δblock_ adjustments (Fig. 3B, right panel), unlike what was observed for %MEP_J8-Leg_Δblock_.

While the prime-mover (i.e. %MEP_J8-FDI_) showed no block- or precue-related modulation, it is important to note that it was globally suppressed at Jump8 relative to baseline (100%), which may have limited sensitivity to detect any broad facilitatory expression of the modulation. Specifically, one-sample t-tests against baseline (100%) revealed significant suppression of %MEP_J8-FDI_ only in the Ctrl_Low_–Rew_High_ and Ctrl_High_-Rew_High_ conditions (all t > -3.52, all p-corrected < 0.008; Fig. 3B, left panel). A comparable pattern of suppression was found for %MEP_J8-FDI_ on the unchosen side; corresponding analyses and the associated figures are reported in the Supplementary Materials (Suppl. Fig. 3A).

##### Surround inhibition

Even with the prime mover suppressed, surround inhibition predicts an additional suppression in neighboring muscles. We quantified this using Δ%MEP_J8-APB_ (%MEP_J8-APB_ minus %MEP_J8-FDI_) on the chosen side, where Δ values below 0% indicate stronger suppression in APB than in FDI. One-sample t-tests showed that Δ%MEP_J8-APB_ was significantly negative only in the Ctrl_Low_–Rew_High_ and Ctrl_High_-Rew_High_ conditions (both t > −3.94, both p-corrected < 0.036; Fig. 3C, left panel). Moreover, Δ%MEP_J8-APB_ showed no effect of *block type* (F_(1, 934)_ = 2.11, p = 0.146), This absence of effect was further confirmed by a Bayesian ANOVA, which provided evidence in favor of the null model (BF_10_ = 0.35), indicating comparable levels of surround inhibition across block types. In contrast, we observed a significant effect of *precue type* (F_(1, 934)_ = 6.95, p = 0.009; Fig. 3C, left panel), with more negative values following Rew_High_ than Rew_Low_ precues. This precue effect further interacted with UPPS_Urgency_ (*precue type × UPPS_Urgency_* interaction: F_(1, 934)_ = 5.56, p = 0.019). Simple-effects analyses at representative UPPS_Urgency_ values (illustrated as Δprecue in Fig. 3C, middle panel) showed a significant Rew_High_–Rew_Low_ difference at low (F_(1, 934)_ = 11.47, p <0.001) and average UPPS_Urgency_ (F_(1, 934)_ = 6.95, p = 0.009), but not at high UPPS_Urgency_ (F_(1, 934)_ = 0.04, p = 0.843). This pattern contrasts with the urgency results, as the largest reward-related increase in surround inhibition occurred in participants who showed the weakest reward-related modulation of urgency. Note that only a main effect of *precue type* (F_(1, 934)_ = 14.68, p < 0.001; see Fig. 3C, right panel), with no interaction with UPPS_Urgency_ (*precue type × UPPS_Urgency_* interaction: F_(1, 934)_ = 1.41, p = 0.235), was found when considering directly %MEP APB values (i.e. %MEP_J8-APB_) rather than the Δ (APB-FDI) values (i.e. Δ%MEP_J8-APB_). Finally, Δ%MEP_J8-APB_ effects were mostly confined to the responding (chosen) side. On the unchosen side, Δ%MEP_J8-APB_ remained close to 0% and showed no consistent suppression across conditions, with no significant main effects or interactions (Supplementary Materials; Suppl. Fig. 3B). This lateralized pattern is consistent with a reward-related modulation selectively targeting motor representations surrounding the engaged effector.

Taken together, these findings suggest that surround inhibition implements reward prospect (but not urgency) in the motor system. Specifically, surround inhibition increased with higher rewards even though higher rewards were associated with down-regulation of urgency. Moreover, surround inhibition varied inversely with the magnitude of individuals’ urgency adjustments.

#### Baseline corticospinal excitability

In a final analysis, we examined whether resting corticospinal excitability measured within blocks (elicited at TMS_Baseline-in_) differed from baseline measured outside the blocks (elicited at TMS_Baseline-out_), and whether this modulation differed between Ctrl_Low_ and Ctrl_High_ blocks. To do so, we expressed MEP amplitudes for leg, FDI and APB MEP amplitudes at TMS_Baseline-in_ (B-in) as a percentage of TMS_Baseline-out_ (%MEP_B-in-Leg_, %MEP_B-in-FDI_ and %MEP_B-in-APB_). For hand muscles, MEPs were pooled across both sides. One-sample t-tests showed that all three baseline probes were significantly greater than 100% (t > 2.74, p-corrected < 0.016; Fig. 3D, left panel) in both Ctrl_Low_ and Ctrl_High_ blocks, indicating overall higher baseline excitability within blocks than outside blocks.

Linear mixed-effects models revealed no main effect of *block type* and no *block type x UPPS_Urgency_* interaction for baseline hand excitability (%MEP_B-in-FDI_ or %MEP_B-in-APB_; all F < 0.96, all p > 0.33; see Fig. 3D, left, right-middle and right-most panels), nor for baseline leg excitability (%MEP_B-in-Leg_; all F < 3.35, all p > 0.074; Fig. 3D, left and left-middle panels).

Together, these null baseline effects indicate that neither block demands nor impulsivity influenced corticospinal excitability prior to deliberation, ruling out baseline differences as a source of the observed effects during the decision period. Instead, the motor signatures reported above reflect dynamic, decision-specific modulations emerging during deliberation, supporting the view that control-based and reward-based urgency regulation operates through phasic motor adjustments rather than sustained baseline changes.

## 4. Discussion

Although prior work suggests that both control demands and motivation by reward can shape urgency, these influences had not previously been dissociated within a single design to characterize them selectively and probe their distinct motor implementation. Here, we show that control- and reward-driven urgency regulation modes are differentially expressed across individuals, with control-driven adjustments most evident in less impulsive participants and reward-driven adjustments most evident in more impulsive participants. Moreover, control-driven urgency regulation was associated with global modulations of motor activity, whereas reward-driven urgency adjustments were associated with changes in surround inhibition.

Our results align with prior TMS work showing robust surround inhibition during action preparation, indexed here by reduced excitability in the thumb muscle surrounding the prime mover (the index finger muscle). This suppression is thought to sharpen the motor command, limit spillover into irrelevant close-by effectors, and has been interpreted as a gain-control mechanism that enhances signal-to-noise by dampening background activity around the active representation (Beck & Hallett, 2011; Bestmann & Duque, 2016; Duque et al., 2017; Leodori et al., 2019; Derosiere et al., 2022; Greenhouse, 2022). Consistent with our hypothesis and prior work linking this motor signature to reward prospect (Tecilla et al., 2022; Derosiere et al., 2025; Thura et al., 2025), surround inhibition primarily tracked reward-driven urgency adjustments: it differed significantly between low and high reward precues. Critically, surround inhibition increased after high-reward precues even if participants concurrently down-regulated urgency to maximize reward attainment. This pattern challenges the idea that surround inhibition provides a direct physiological readout of urgency (Derosiere et al., 2022) and helps resolve an ambiguity in earlier work in which higher reward co-occurred with higher urgency, making it difficult to disentangle reward-related and urgency-related influences on surround inhibition. By showing that surround inhibition scales with reward prospect even when urgency is strategically reduced, our results instead suggest that surround inhibition is more directly tied to reward value than to urgency per se. This interpretation is consistent with evidence linking surround inhibition to dopaminergic state and motivational invigoration (Tecilla et al., 2022; Derosiere et al., 2025; Thura et al., 2025), including findings that reduced dopamine in Parkinson’s disease is associated with reduced surround inhibition (Jahanshahi & Rothwell, 2017; Wilhelm et al., 2022; Wilhelm et al., 2024). Accordingly, reward-related increases in surround inhibition may still reflect an invigoration motor gain signal, but their behavioral expression can be counteracted by concurrent goal-driven mechanisms that deliberately slow commitment to protect accuracy under high stakes. Notably, the reward-related deepening of surround inhibition was strongest in less impulsive participants, who showed the smallest reward-driven slowing, whereas more impulsive participants showed the largest reward-driven slowing but a weaker deepening of surround inhibition following high reward precues. This pattern suggests that, in more impulsive individuals, reward may still engage invigoration-related motor signals, but that the dominant reward-driven regulation mode involves actively suppressing this invigoration to support cautious, accuracy-oriented behavior, resulting in a reduced strengthening of surround inhibition under high reward in the more impulsive participants.

As predicted, broad modulation appears to be more directly involved in implementing control-driven urgency adjustments. Broad modulation, indexed here by corticospinal excitability in task-irrelevant leg muscles, took the form of a general facilitation, and this facilitation was attenuated when urgency was down-regulated in high control relative to low control blocks, consistent with the idea that adopting a more cautious policy involves a global dampening of motor readiness across the motor system (Klein et al., 2014; Derosiere et al., 2018; Derosiere et al., 2020; Quoilin et al., 2020). This relationship also emerged across individuals: participants showing the strongest control-driven reduction in broad facilitation exhibited the largest decreases in urgency. Mechanistically, this broad modulation likely reflects the balance between non-specific arousal-related upregulation of corticospinal excitability during task engagement (Bundt et al., 2019; Valappil et al., 2025; Weijs et al., 2025) and top-down control processes that dampen global baseline readiness under higher demands, consistent with accounts linking inhibitory control to widespread (non–muscle-specific) changes in corticospinal excitability (Quoilin & Derosiere, 2015; Iacullo et al., 2020). In an accumulation-to-threshold framework, baseline motor readiness can be viewed as setting the starting state of the response process: higher readiness places the system closer to the decision bound, whereas dampening readiness increases the distance to threshold and requires more build-up before the bound is crossed (Wiecki & Frank, 2013; Steinemann et al., 2018; Yau et al., 2020). Notably, reward-driven urgency adjustments were not accompanied by corresponding changes in broad modulation: the broad facilitation pattern did not show attenuation following high versus low reward precues despite the concurrent down-regulation of urgency, indicating that urgency can be regulated without necessarily being implemented through changes in broad facilitation. One possibility is that high reward precues simultaneously enhance arousal-related global facilitation while also recruiting stronger inhibitory control to support caution (Suzuki et al., 2019; Thura, 2020), yielding opposing influences that cancel out at the level of broad facilitation. Together with the surround inhibition findings, this supports a nuanced view in which broad modulation implements global motor readiness and thus preferentially supports control-driven urgency regulation. In contrast, reward-driven urgency adjustments may involve a combination of gain/invigoration and compensatory top-down control, such that this type of urgency change does not necessarily translate into detectable changes in broad modulation.

In principle, if broad modulation reflects a global change in motor readiness, it should not be confined to distant muscles such as the leg: one might also expect some signature in the index finger muscle, especially in less impulsive participants where goal-driven urgency regulation was most evident. Yet, when we probed the index finger muscle at Jump8, we observed no modulation by block or precue, and responses were overall below baseline, which likely reduced our ability to detect any broad facilitatory effect in a task-relevant hand muscle at that time point. This pattern fits well with the extensive literature on preparatory suppression, which repeatedly reports reduced excitability in response-related hand muscles during action preparation and selection (Duque et al., 2017; Hannah et al., 2018; Ibáñez et al., 2020). This literature has proposed that such suppression helps to hold back premature actions (Grandjean & Duque, 2020) but may also set favorable conditions for a sharper subsequent release of the selected response, effectively increasing the contrast between relevant and irrelevant motor activity (Greenhouse et al., 2015). Within this framework, the suppression we observed in the index finger muscle may reflect a broadly “held-back” state of the response system during deliberation, as described in the preparatory suppression literature. At the same time, surround inhibition is commonly interpreted as a way to increase contrast around the selected effector by limiting recruitment of nearby, irrelevant muscles. Rather than implying two fully distinct processes, it is more likely that these influences overlap and jointly contribute to the pattern observed at our Jump8 probe: a generally reduced excitability in the index finger muscle, together with additional suppression in surrounding hand representations, here indexed by the thumb muscle.

A final point concerns the behavioral expression of the control-driven and reward-driven urgency adjustments and their relationship to impulsivity. The performance data revealed that participants implemented control-driven adjustments by slowing their decisions and improving accuracy when caution was required. These adjustments varied systematically with impulsivity. Less impulsive individuals showed the strongest block-related slowing, consistent with their larger down regulation of urgency, and with evidence that more impulsive individuals exhibit diminished ability to increase caution when task demands require it (Bari & Robbins, 2013). In contrast, reward-driven adjustments were more pronounced in more impulsive participants, who exhibited robust slowing following high reward precues, translating into improved accuracy. This finding is consistent with evidence that impulsivity is tightly linked to reward sensitivity and reward–driven adjustments in decision policy (Dalley & Robbins, 2017; Jauregi et al., 2018). Together, these findings align with models proposing that individuals differ in how they regulate urgency and decision policies across contexts, including variations tied to trait-like impulsivity (Thura et al., 2025).

Our results show that the impact of control demands and reward prospect can be dissociated as two modes of regulation, and that their relative expression varies across individuals: less impulsive participants primarily adjusted urgency to task control demands, whereas more impulsive participants showed stronger reward-related urgency adjustments. This pattern indicates that impulsivity does not simply reflect “more urgency” but a different weighting of the signals that shape it. Motor-level probes revealed how urgency regulation is instantiated within the motor system. Broad modulation implemented the control-driven mode: motor activity was more strongly attenuated when participants adopted a cautious policy, an effect that was clearest in less impulsive participants. By contrast, surround inhibition implemented the reward-driven mode, consistent with a motor invigoration mechanism that can increase with reward even when urgency is strategically down-regulated. The weaker reward-related strengthening of surround inhibition in more impulsive participants suggests that reward-driven urgency regulation may involve counteracting this invigoration. Together, these findings show that urgency arises from the interaction of distinct control- and reward-driven processes that are differentially implemented within the motor system, and that impulsivity biases which of these influences predominates.

## Author contributions

T.F., F.C., F.F., J.L., P.V., G.D., and J.D. designed research; T.F., F.C., A.S., F.F. performed researd; T.F., F.C., A.S., J.L., and J.D. analyzed data; T.F., F.C., and J.D. wrote the first draft of the paper; T.F., F.C., A.S., F.F., J.L., P.V., G.D., and J.D edited the paper; T.F., F.C., and J.D. wrote the paper.

## Acknowledgements

T.F. was supported by the Fund for Research training in Industry and Agriculture (FRIA/FNRS: FC49813) and by UCLouvain (FSR: 01146594). F.C. was supported by the Fund for Research training in Industry and Agriculture (FRIA/FNRS: FC60653) and by UCLouvain (FSR: 01147926). The work was also supported by grants from the Belgian National Fund for Scientific Research (PDR UrgeToAct-40013512) and from the UCLouvain (ARC Coaction).

## References

1. Barakat N, Brunelin J, Cailhol L, Saint-Amant A, Paoletti LM, Poulet E, Neige C, & Vallet W. (2025). Inhibitory control deficits in borderline personality disorder: a meta-analysis of stop-signal and Go/No-Go tasks. Psychol Med, 55, e319.

2. Bari A, & Robbins TW. (2013). Inhibition and impulsivity: Behavioral and neural basis of response control. Progress in Neurobiology, 108, 44–79.

3. Bauer DJ, & Curran PJ. (2005). Probing Interactions in Fixed and Multilevel Regression: Inferential and Graphical Techniques. Multivariate Behav Res, 40(3), 373–400.

4. Beck S, & Hallett M. (2011). Surround inhibition in the motor system. Exp Brain Res, 210(2), 165–172.

5. Bestmann S, & Duque J. (2016). Transcranial Magnetic Stimulation:Decomposing the Processes Underlying Action Preparation. The Neuroscientist, 22(4), 392–405.

6. Bundt C, Bardi L, Verbruggen F, Boehler CN, Brass M, & Notebaert W. (2019). Reward anticipation changes corticospinal excitability during task preparation depending on response requirements and time pressure. Cortex, 120, 159–168.

7. Carland MA, Thura D, & Cisek P. (2019). The Urge to Decide and Act: Implications for Brain Function and Dysfunction. The Neuroscientist, 25(5), 491–511.

8. Carsten T, Fievez F, & Duque J. (2023). Movement characteristics impact decision-making and vice versa. Sci Rep, 13(1), 3281.

9. Cisek P, Puskas GA, & El-Murr S. (2009). Decisions in changing conditions: the urgency-gating model. J Neurosci, 29(37), 11560–11571.

10. Cyders MA, Littlefield AK, Coffey S, & Karyadi KA. (2014). Examination of a short English version of the UPPS-P Impulsive Behavior Scale. Addict Behav, 39(9), 1372–1376.

11. Dalley J, & Robbins T. (2017). Fractionating impulsivity: Neuropsychiatric implications. Nature Reviews Neuroscience, 18, 158–171.

12. Deng ZD, Lisanby SH, & Peterchev AV. (2014). Coil design considerations for deep transcranial magnetic stimulation. Clin Neurophysiol, 125(6), 1202–1212.

13. Derosiere G, Klein P-A, Nozaradan S, Zénon A, Mouraux A, & Duque J. (2018). Visuomotor correlates of conflict expectation in the context of motor decisions. Journal of Neuroscience, 38(44), 9486–9504.

14. Derosiere G, Shokur S, & Vassiliadis P. (2025). Reward signals in the motor cortex: from biology to neurotechnology. Nat Commun, 16(1), 1307.

15. Derosiere G, Thura D, Cisek P, & Duque J. (2019). Motor cortex disruption delays motor processes but not deliberation about action choices. Journal of neurophysiology, 122(4), 1566–1577.

16. Derosiere G, Thura D, Cisek P, & Duque J. (2021). Trading accuracy for speed over the course of a decision. Journal of Neurophysiology, 126(2), 361–372.

17. Derosiere G, Thura D, Cisek P, & Duque J. (2022). Hasty sensorimotor decisions rely on an overlap of broad and selective changes in motor activity. Plos Biology, 20(4), e3001598.

18. Derosiere G, Vassiliadis P, & Duque J. (2020). Advanced TMS approaches to probe corticospinal excitability during action preparation. Neuroimage, 213, 116746.

19. Drew DS, Muhammed K, Baig F, Kelly M, Saleh Y, Sarangmat N, Okai D, Hu M, Manohar S, & Husain M. (2020). Dopamine and reward hypersensitivity in Parkinson’s disease with impulse control disorder. Brain, 143(8), 2502–2518.

20. Duque J, Greenhouse I, Labruna L, & Ivry RB. (2017). Physiological markers of motor inhibition during human behavior. Trends in neurosciences, 40(4), 219–236.

21. Fievez F, Cos I, Carsten T, Derosiere G, Zénon A, & Duque J. (2024). Task goals shape the relationship between decision and movement speed. J Neurophysiol, 132(6), 1837–1856.

22. Frömer R, Lin H, Dean Wolf CK, Inzlicht M, & Shenhav A. (2021). Expectations of reward and efficacy guide cognitive control allocation. Nat Commun, 12(1), 1030.

23. Grandjean J, Derosiere G, Vassiliadis P, Quemener L, de Wilde Y, & Duque J. (2018). Towards assessing corticospinal excitability bilaterally: Validation of a double-coil TMS method. Journal of Neuroscience Methods, 293, 162–168.

24. Grandjean J, & Duque J. (2020). A TMS study of preparatory suppression in binge drinkers. Neuroimage Clin, 28, 102383.

25. Greenhouse I. (2022). Inhibition for gain modulation in the motor system. Experimental Brain Research, 240(5), 1295–1302.

26. Greenhouse I, Sias A, Labruna L, & Ivry RB. (2015). Nonspecific inhibition of the motor system during response preparation. Journal of Neuroscience, 35(30), 10675–10684.

27. Hannah R, Cavanagh SE, Tremblay S, Simeoni S, & Rothwell JC. (2018). Selective Suppression of Local Interneuron Circuits in Human Motor Cortex Contributes to Movement Preparation. J Neurosci, 38(5), 1264–1276.

28. Iacullo C, Diesburg DA, & Wessel JR. (2020). Non-selective inhibition of the motor system following unexpected and expected infrequent events. Exp Brain Res, 238(12), 2701–2710.

29. Ibáñez J, Hannah R, Rocchi L, & Rothwell JC. (2020). Premovement Suppression of Corticospinal Excitability may be a Necessary Part of Movement Preparation. Cereb Cortex, 30(5), 2910–2923.

30. Jahanshahi M, & Rothwell JC. (2017). Inhibitory dysfunction contributes to some of the motor and non-motor symptoms of movement disorders and psychiatric disorders. Philos Trans R Soc Lond B Biol Sci, 372(1718).

31. Jauregi A, Kessler K, & Hassel S. (2018). Linking Cognitive Measures of Response Inhibition and Reward Sensitivity to Trait Impulsivity. Front Psychol, 9, 2306.

32. Klein PA, Petitjean C, Olivier E, & Duque J. (2014). Top-down suppression of incompatible motor activations during response selection under conflict. Neuroimage, 86, 138–149.

33. Leodori G, Thirugnanasambandam N, Conn H, Popa T, Berardelli A, & Hallett M. (2019). Intracortical Inhibition and Surround Inhibition in the Motor Cortex: A TMS-EEG Study. Front Neurosci, 13, 612.

34. Manohar S, Chong T, Apps M, Batla A, Stamelou M, Jarman P, Bhatia K, & Husain M. (2015). Reward Pays the Cost of Noise Reduction in Motor and Cognitive Control. Current biology : CB, 25.

35. Murphy J, Devue C, Corballis PM, & Grimshaw GM. (2020). Proactive Control of Emotional Distraction: Evidence From EEG Alpha Suppression. Front Hum Neurosci, 14, 318.

36. Niehaus L, Meyer BU, & Weyh T. (2000). Influence of pulse configuration and direction of coil current on excitatory effects of magnetic motor cortex and nerve stimulation. Clin Neurophysiol, 111(1), 75–80.

37. Oldfield RC. (1971). The assessment and analysis of handedness: The Edinburgh inventory. Neuropsychologia, 9(1), 97–113.

38. Quoilin C, & Derosiere G. (2015). Global and Specific Motor Inhibitory Mechanisms during Action Preparation. J Neurosci, 35(50), 16297–16299.

39. Quoilin C, Fievez F, & Duque J. (2019). Preparatory inhibition: Impact of choice in reaction time tasks. Neuropsychologia, 129, 212–222.

40. Quoilin C, Grandjean J, & Duque J. (2020). Considering Motor Excitability During Action Preparation in Gambling Disorder: A Transcranial Magnetic Stimulation Study. Front Psychiatry, 11, 639.

41. Reppert TR, Heitz RP, & Schall JD. (2023). Neural mechanisms for executive control of speed-accuracy trade-off. Cell Rep, 42(11), 113422.

42. Rossini PM, et al. (2015). Non-invasive electrical and magnetic stimulation of the brain, spinal cord, roots and peripheral nerves: Basic principles and procedures for routine clinical and research application. An updated report from an I.F.C.N. Committee. Clin Neurophysiol, 126(6), 1071–1107.

43. Steinemann NA, O’Connell RG, & Kelly SP. (2018). Decisions are expedited through multiple neural adjustments spanning the sensorimotor hierarchy. Nat Commun, 9(1), 3627.

44. Suzuki M, Suzuki T, Wang YJ, & Hamaguchi T. (2019). Changes in Magnitude and Variability of Corticospinal Excitability During Rewarded Time-Sensitive Behavior. Front Behav Neurosci, 13, 147.

45. Tecilla M, Guerra A, Rocchi L, Määttä S, Bologna M, Herrojo Ruiz M, Biundo R, Antonini A, & Ferreri F. (2022). Action selection and motor decision making: insights from transcranial magnetic stimulation. Brain Sciences, 12(5), 639.

46. Thirugnanasambandam N, Leodori G, Popa T, Kassavetis P, Mandel A, Shaft A, Kee J, Kashyap S, Khodorov G, & Hallett M. (2020). Parietal conditioning enhances motor surround inhibition. Brain Stimul, 13(2), 447–449.

47. Thura D. (2020). Decision urgency invigorates movement in humans. Behavioural Brain Research, 382, 112477.

48. Thura D, Cabana JF, Feghaly A, & Cisek P. (2022). Integrated neural dynamics of sensorimotor decisions and actions. PLoS Biol, 20(12), e3001861.

49. Thura D, & Cisek P. (2014). Deliberation and Commitment in the Premotor and Primary Motor Cortex during Dynamic Decision Making. Neuron, 81(6), 1401–1416.

50. Thura D, Haith AM, Derosiere G, & Duque J. (2025). The integrated control of decision and movement vigor. Trends Cogn Sci, 29(12), 1146–1157.

51. Valappil AC, Grilc N, Castelli F, Chye S, Wright DJ, Tyler CJ, Knight R, Mian OS, Tillin NA, & Bruton AM. (2025). Corticospinal excitability is facilitated during coordinative action observation and motor imagery. Cerebral Cortex, 35(6).

52. Vassiliadis P, Derosiere G, Grandjean J, & Duque J. (2020). Motor training strengthens corticospinal suppression during movement preparation. Journal of Neurophysiology, 124(6), 1656–1666.

53. Vassiliadis P, Grandjean J, Derosiere G, de Wilde Y, Quemener L, & Duque J. (2018). Using a Double-Coil TMS Protocol to Assess Preparatory Inhibition Bilaterally. Front Neurosci, 12, 139.

54. Weijs ML, Missura S, Potok-Szybińska W, Bächinger M, Badii B, Carro-Domínguez M, Wenderoth N, & Meissner SN. (2025). Modulating cortical excitability and cortical arousal by pupil self-regulation. Nat Commun, 16(1), 4552.

55. Wiecki TV, & Frank MJ. (2013). A computational model of inhibitory control in frontal cortex and basal ganglia. Psychol Rev, 120(2), 329–355.

56. Wilhelm E, Derosiere G, Quoilin C, Cakiroglu I, Paço S, Raftopoulos C, Nuttin B, & Duque J. (2024). Subthalamic DBS does not restore deficits in corticospinal suppression during movement preparation in Parkinson’s disease. Clin Neurophysiol, 165, 107–116.

57. Wilhelm E, Quoilin C, Derosiere G, Paço S, Jeanjean A, & Duque J. (2022). Corticospinal suppression underlying intact movement preparation fades in late Parkinson’s disease. *medRxiv*, 2022.2002.2003.22269055.

58. Yadav G, Vassiliadis P, Dubuc C, Hummel FC, Derosiere G, & Duque J. (2025). Effect of Extrinsic Reward on Motor Plasticity during Skill Learning. eNeuro, 12(4).

59. Yau Y, Dadar M, Taylor M, Zeighami Y, Fellows LK, Cisek P, & Dagher A. (2020). Neural Correlates of Evidence and Urgency During Human Perceptual Decision-Making in Dynamically Changing Conditions. Cereb Cortex, 30(10), 5471–5483.

60. Yau Y, Hinault T, Taylor M, Cisek P, Fellows LK, & Dagher A. (2021). Evidence and urgency related EEG signals during dynamic decision-making in humans. Journal of Neuroscience, 41(26), 5711–5722.

