## Supplementary Materials for "Motor implementation of control and reward-based urgency regulation across impulsivity"

*Shared contribution

¥Corresponding author

### Supplementary materials

#### 2. Materials and Methods

##### 2.5 Data preprocessing and endpoint measures

***Behavior in the Tokens task: urgency and performance-related endpoint measures***

*Justification for pooling trials across TMS conditions and blocks with sessions*

Behavioral analyses reported in the main text were run regardless of TMS conditions (no stimulation, $TMS_{Baseline\text{-}in}$, or $TMS_{Jump8}$). This is because preliminary analyses confirmed that decision duration and accuracy were comparable across conditions (see Suppl. Fig. 1A for decision duration, left panel). Moreover, to account for potential learning effects within and across sessions, we fitted an additional model including block number and session as fixed effects. Decisions were generally faster in the $TMS_{Leg}$ session (Suppl. Fig. 1A, middle panel), but this did not affect our main analyses because behavioral and motor excitability measures were never compared across sessions, and session order was fixed. Finally, decision durations remained stable across blocks except for the first block, which showed shorter decision durations (Suppl. Fig. 1A, right panel). Analyses conducted with and without this block yielded similar results; therefore, all trials were retained.

***TMS probes in the Tokens task: broad modulation and surround inhibition***

*RT-matching procedure for MEP selection at $TMS_{Jump8}$*

To eliminate variability in MEP amplitude that could arise from differences in reaction time (RT) across trials or conditions, all analyses of MEPs elicited at $TMS_{Jump8}$ (1400 ms) were performed on a strict RT-matched subset of trials, following the procedure described in the main text.

As in the main text, RT matching was based on a single automatically selected window. Candidate RT windows (100-ms steps) were evaluated within a predefined RT range, extending from 200 ms after the TMSJump8 pulse (i.e., ≥1600 ms, to avoid contamination of MEPs by movement execution) to the end of the trial (2800 ms). A window was considered valid if every subject × condition cell contained at least 12 trials, allowing trial selection to be aligned on the least represented condition. Among valid candidates, the narrowest window was selected to minimize RT-related confounds; when several windows met this criterion, the window closest to the TMSJump8 pulse (1400 ms) was chosen.

Using this procedure, the selected RT window was [1900–2400 ms] for the $\text{TMS}_{\text{Hand}}$ session and [1700–2400 ms] for the $\text{TMS}_{\text{Leg}}$ session. In both sessions, aligning trial counts on the least frequent condition resulted in the inclusion of exactly 12 trials per subject and per condition, ensuring that all subjects contributed equally to group-level analyses. When more than 12 trials were available within the selected window, trials were selected deterministically, prioritizing those with RTs closest to the condition-specific median RT. For the $\text{TMS}_{\text{Leg}}$ session, an additional constraint enforced a balanced number of left and right choices within each condition (≥6 per side), as described in the main text.

Before RT matching, median RTs differed across conditions within each session (Suppl. Fig. 1B, left panels). In the $\text{TMS}_{\text{Hand}}$ session, median RTs equaled 2125 ± 15 ms [$\text{Ctrl}_{\text{Low}}$–$\text{Rew}_{\text{Low}}$], 2165 ± 13 ms [$\text{Ctrl}_{\text{Low}}$–$\text{Rew}_{\text{High}}$], 2177 ± 13 ms [$\text{Ctrl}_{\text{High}}$–$\text{Rew}_{\text{Low}}$], and 2212 ± 12 ms [$\text{Ctrl}_{\text{High}}$–$\text{Rew}_{\text{High}}$]. In the $\text{TMS}_{\text{Leg}}$ session, median RTs before matching were also disparate, including 2097 ± 11 ms [$\text{Ctrl}_{\text{Low}}$–$\text{Rew}_{\text{Low}}$], 2132 ± 11 ms [$\text{Ctrl}_{\text{Low}}$–$\text{Rew}_{\text{High}}$], 2097 ± 10 ms [$\text{Ctrl}_{\text{High}}$–$\text{Rew}_{\text{Low}}$], and 2157 ± 13 ms [$\text{Ctrl}_{\text{High}}$–$\text{Rew}_{\text{High}}$].

After RT matching, RT distributions strongly overlapped across conditions (Suppl. Fig. 1B, right panels), with median RTs converging within a comparable range. Post-matching median RTs equaled, for the $\text{TMS}_{\text{Hand}}$ session: 2145 ± 7 ms [$\text{Ctrl}_{\text{Low}}$–$\text{Rew}_{\text{Low}}$],: 2151 ± 7 ms [$\text{Ctrl}_{\text{Low}}$–$\text{Rew}_{\text{High}}$], 2155 ± 6 ms [$\text{Ctrl}_{\text{High}}$–$\text{Rew}_{\text{Low}}$], and 2142 ± 7 ms [$\text{Ctrl}_{\text{High}}$–$\text{Rew}_{\text{High}}$]. Then for the

$TMS_{Leg}$ session, matched RTs equaled : 2047 ± 6 ms [$Ctrl_{Low}$–$Rew_{Low}$], 2042 ± 7 ms [$Ctrl_{Low}$–$Rew_{High}$], 2047 ± 6 ms [$Ctrl_{High}$–$Rew_{Low}$], 2052 ± 6 ms [$Ctrl_{High}$–$Rew_{High}$].

Together, this procedure ensured that $TMS_{Jump8}$ MEPs were compared within a matched and standardized RT range across all conditions, thereby eliminating RT-related confounds while maintaining balanced datasets with identical trial counts across subjects.

#### 3. Results

##### *Behavior in the Tokens task: urgency and performance-related endpoint measures*

*Urgency-related endpoint measures: U-Intercept and U-Slope*

Because U-Intercept (initial level) and U-Slope (growth rate) are estimated from the same linear function and share variance, we isolated their unique contributions via residualization. Specifically, we regressed Intercept on Slope and Slope on Intercept, and used the residuals as orthogonalized indices capturing variance not shared with the companion parameter.

Regarding U-Intercept, which reflects initial urgency, the linear mixed-effects analysis revealed no main effect of *block type* ($F_{(1, 54)} = 1.77$, $p = 0.189$; Suppl. Fig. 2A, left-most panel), but a significant *block type x* $UPPS_{Urgency}$ interaction ($F_{(1, 54)} = 4.32$, $p = 0.042$), indicating that blockrelated adjustments of U-Intercept depended on impulsivity. To probe this interaction, we ran a post-hoc analysis examining the simple effects of *block type* on U-Intercept (illustrated as a delta [Δblock: $Ctrl_{High}$ – $Ctrl_{Low}$] in Suppl. Fig. 2A, left-middle panel) at three representative $UPPS_{Urgency}$ values, with "low" corresponding to the mean $UPPS_{Urgency}$ minus 1 SD, "average" corresponding to the mean $UPPS_{Urgency}$, and "high" corresponding to the mean $UPPS_{Urgency}$ plus 1 SD. At low $UPPS_{Urgency}$ values, U-Intercept was lower in $Ctrl_{High}$ than $Ctrl_{Low}$ blocks ($F(1, 54) = 5.82$, $p = 0.019$), consistent with a down-regulation of urgency in $Ctrl_{High}$ blocks among less impulsive participants. This slowing was not observed at average or high $UPPS_{Urgency}$ values (all $F < 1.77$, all $p > 0.189$). This heterogeneity likely accounts for the absence of a *block type* effect at the group level. The U-Intercept was also influenced by whether participants would be rewarded for a correct response on a given trial, as indicated by a significant main effect of *precue type* ($F_{(1, 54)} = 12.44$, $p < 0.001$; Suppl. Fig. 2A, right-middle panel): U-Intercept was lower in $Rew_{High}$ trials (-0.005 ± 0.007 a.u.) than in $Rew_{Low}$ trials (0.005 ± 0.007 a.u.). This effect interacted with $UPPS_{Urgency}$ ($F_{(1, 54)} = 6.31$, $p = 0.015$). Post-hoc simple-effect tests of *precue type* (illustrated as a delta [Δprecue: $Rew_{High}$ minus $Rew_{Low}$] in Suppl. Fig. 2A, right-most panel) at representative $UPPS_{Urgency}$ values showed a significant *precue type* effect at average and high $UPPS_{Urgency}$ scores (both $F_{(1, 54)} > 12.44$, both $p < 0.001$) but not at the lowest scores ($F_{(1, 54)} = 0.50$, $p = 0.484$). No additional interactions were observed on Intercept (all $F < 1.78$, all $p > 0.188$). Note that all these effects are similar to those reported for U-Jump8 in the main manuscript.

Regarding U-Slope, which reflects the rate at which urgency rises over the course of a trial, there was a main effect of *block type* ($F_{(1,54)} = 5.64$, $p = 0.021$; Suppl. Fig. 2B, left-most panel), with trials displaying a higher U-Slope in $Ctrl_{High}$ than $Ctrl_{Low}$ block. This effect did not show any interaction with $UPPS_{Urgency}$ ($F_{(1,54)} = 0.06$, $p = 0.803$), as evident from the consistent Δblock values across low, average and high $UPPS_{Urgency}$ values (Suppl. Fig. 2B, left-middle panel). There was also a main effect of *precue type* ($F_{(1, 54)} = 7.67$, $p = 0.008$; Suppl. Fig. 2B, right-middle panel), with higher U-Slope following $Rew_{High}$ than $Rew_{Low}$ precues, but again the *precue type x* $UPPS_{Urgency}$ interaction was not significant ($F_{(1,54)} = 3.88$, $p = 0.054$), as evident from the consistent Δprecue values across low, average and high $UPPS_{Urgency}$ values (see Suppl. Fig. 2B, right-most panel). No additional interactions were observed on U-Slope (all $F < 3.76$, all $p > 0.058$). Hence, U-Slope was generally higher in $Ctrl_{High}$ blocks and following $Rew_{High}$ precues.

*Performance-related endpoint measures: Additional information on trial type effects*

Starting with decision durations, the linear mixed-effects analysis revealed a main effect of *trial type* ($F_{(2,\ 23083)}$ = 5577.13, p < 0.001). Decisions were slower in ambiguous than in both misleading trials (post-hoc Tukey tests z = 36.8, p < 0.001) and obvious trials (z = 105.3, p < 0.001), with decisions in misleading trials also found slower than in obvious ones (z = –55.6, p < 0.001), as expected based on the different token dynamics. As briefly mentioned in the main text (and shown on Fig. 2C, left-most panel), the *trial type* × *block type* interaction was also significant ($F_{(2,\ 23083)}$ = 35.12, p < 0.001), with post-hoc analyses revealing a significant effect of *block type* within all three trial types (all z > –13.49, all p < 0.001). This suggests that the slower decision durations observed in $Ctrl_{High}$ blocks cannot be solely explained by the higher proportion of misleading trials. Instead, participants appear to have adopted a generally slower decision strategy to better cope with the increased likelihood of misleading evidence, in line with the effects observed on urgency parameters.

Regarding accuracy, the main effect of *trial type* suggest that it was globally lower in misleading than ambiguous trials ($\chi^2(1)$ = 1633,50, p < 0.001). Yet, as briefly mentioned in the main text (and shown in Fig. 2D, left-most panel), the *trial type* × *block type* interaction was also significant ($\chi^2_{(1)}$ = 11.17, p < 0.001). Post-hoc comparisons revealed that accuracy was higher for misleading trials in $Ctrl_{High}$ compared with $Ctrl_{Low}$ blocks (z = –5.57, p < 0.001), consistent with the idea that participants benefited from the slower decision policy in this context. In contrast, no accuracy gain was observed for ambiguous trials (z = –1.27, p = 0.585), suggesting that the additional decision time in $Ctrl_{High}$ blocks was insufficient to substantially increase the amount of available evidence in these trials. Thus, while participants adapted their decision speed to the increased uncertainty of $Ctrl_{High}$ blocks (i.e. consistent with the lower U-Intercept reported in the previous section and the lower U-Jump8 in the main text), this adjustment proved beneficial only when the misleading evidence could be resolved through additional evidence accumulation.

***TMS probes in the Tokens task: broad modulation and Surround inhibition***

*Broad modulation*

The main-text analyses on $\%MEP_{J8\text{-}FDI}$ focused on the probe recorded on the chosen side (i.e., right $\%MEP_{J8\text{-}FDI}$ for right-hand trials and left $\%MEP_{J8\text{-}FDI}$ for left-hand trials). This choice was motivated by our aim to test for facilitation of the prime mover; however, $\%MEP_{J8\text{-}FDI}$ on the chosen side was globally below baseline (see main text Results and Fig. 3B, left panel). Here, we report complementary analyses for $\%MEP_{J8\text{-}FDI}$ on the unchosen side (left $\%MEP_{J8\text{-}FDI}$ in right-side trials, and right $\%MEP_{J8\text{-}FDI}$ in left-side trials). One-sample t-tests against baseline (100%) showed significant suppression in all conditions (all $t > -3.14$, all p-corrected $< 0.048$), except in the $Ctrl_{High}$ -$Rew_{High}$ condition ($t(19) = -1.77$, p-corrected = 0.184; see Suppl. Fig. 3A, left panel). Moreover, as on the chosen side, % $MEP_{J8\text{-}FDI}$ on the unchosen side did not vary with *block type* or *precue type* (all $F < 0.17$, all $p > 0.679$), and no significant interactions involving *block type, precue type* or $UPPS_{Urgency}$ were found (all $F < 1.85$, all $p > 0.174$).

*Surround inhibition*

Here too, analyses in the main text were restricted to the chosen side, because surround inhibition is thought to operate around the prime mover. To further test the specificity of the observed effects on $\Delta\%MEP_{J8\text{-}APB}$, we additionally analyzed $\Delta\%MEP_{J8\text{-}APB}$ on the unchosen side. One-sample t-tests indicate that $\Delta\%MEP_{J8\text{-}APB}$ was significantly smaller than 0% only in the $Ctrl_{Low}$ -$Rew_{High}$ condition ($t(19) = -2.83$, p-corrected = 0.02; see Suppl. Fig. 3B, left panel). Contrary to findings on the chosen side, there was no *precue type x* $UPPS_{Urgency}$ interaction ($F_{(1, 934)} = 0.63$, $p = 0.428$; Suppl. Fig. 3B, middle panel) and this absence was also true when considering the $\%MEP_{J8\text{-}APB}$ directly rather than the APB-FDI Δ (Suppl. Fig. 3B, right panel). In fact, none of the main effects or interactions involving *block type, precue type* or $UPPS_{Urgency}$ were significant for these two measures on the unchosen side (all $F < 3.86$, all $p > 0.05$).

#### Figure legends

**Supplementary Figure 1. Additional details on data preprocessing procedures.**
**(A) Decision durations across TMS conditions, blocks and sessions.** Bold horizontal lines represent estimated marginal means, error bars indicate standard error, gray lines connect within-participant values, and dots represent individual participants. **Left panel:** Decision durations were comparable in no stimulation, $TMS_{Baseline\text{-}in}$ and $TMS_{Jump8}$ trials. **Middle panel:** Decision durations were generally shorter in the $TMS_{Leg}$ session. **Right panel:** Decision durations were generally shorter in the first block of sessions. **(B) RT matching procedure for MEP selection at $TMS_{Jump8}$.** Distributions of trials as a function of RTs are shown for trials in $Ctrl_{Low}$ (yellow) and $Ctrl_{High}$ (blue) blocks involving $Rew_{Low}$ and $Rew_{High}$ precues (light and dark colors, respectively). Vertical dotted lines represent the median RT for each condition. **Left panel:** Before the matching procedure, distributions were quite different across conditions whether considering the $TMS_{Hand}$ session (on the left) or the $TMS_{Leg}$ session (on the right). **Right panel:** The matching procedure allowed to obtain highly overlapping profiles, indicating that the trial-selection procedure effectively matched RTs across conditions in both the $TMS_{Hand}$ session (on the left) and the $TMS_{Leg}$ session (on the right). ***: p-value < 0.01; ***: p-value < 0.001*

**Supplementary Figure 2. Additional results on (A) U-Intercept and (B) U-slope. Left-most panel:** Urgency parameters are shown as a function of *block type* with $Ctrl_{Low}$ blocks in yellow and $Ctrl_{High}$ blocks in blue. **Left-middle panel:** Urgency parameters are displayed as a delta (Δblock: $Ctrl_{High} - Ctrl_{Low}$) for low, average and high representative $UPPS_{Urgency}$ values. **Right-middle panel:** Urgency parameters are shown as a function of *precue type* with $Rew_{Low}$ in light and $Rew_{High}$ in dark. **Right-most panel:** Urgency parameters are displayed as a delta (Δprecue: $Rew_{High} - Rew_{Low}$) for low, average and high representative $UPPS_{Urgency}$ values. *Bold horizontal lines represent estimated marginal means, error bars indicate standard error, gray lines connect within-participant values, and dots represent individual participants. *: p-value < 0.05; **: p-value < 0.01; ***: p-value < 0.001*

**Supplementary Figure 3. Additional results on TMS probes.**
**(A) $\%MEP_{J8\text{-}FDI}$ on unchosen side. Left panel:** Results are expressed in percentage of baseline, with one-sample t-test comparisons to 100% indicated by # symbols. **Middle panel:** MEPs are presented as a delta (Δblock: $Ctrl_{High} - Ctrl_{Low}$) for low, average and high representative $UPPS_{Urgency}$ values. **Right panel:** MEPs are presented as a delta (Δblock = $Ctrl_{High} - Ctrl_{Low}$) as a function of corresponding change in block type for urgency at Jump8 (Δblock = $Ctrl_{High} - Ctrl_{Low}$). **(B) $\Delta\%MEP_{J8\text{-}APB}$ on unchosen side. Left panel**: Results for $\Delta\%MEP_{J8\text{-}APB}$ (APB – FDI) are presented with comparison one-sample t-test comparison to 0, with negative values indicating smaller MEPs in APB than FDI, indicated by # symbols. **Middle panel:** $\Delta\%MEP_{J8\text{-}APB}$ (APB – FDI) is presented as a delta (Δprecue: $Rew_{High} - Rew_{Low}$) for low, average and high representative $UPPS_{Urgency}$ values. **Right panel:** $\%MEP_{J8\text{-}APB}$ (without FDI) are expressed in percentage of baseline, with one-sample t-test comparison to 100% indicated by # symbols. *Bold horizontal lines represent estimated marginal means, error bars indicate standard error, gray lines connect within-participant values, and dots represent individual participants. #: p-value < 0.05 for Student's t-test*
