## Supplementary figures and images for "Motor implementation of control and reward-based urgency regulation across impulsivity"

### Figure S1

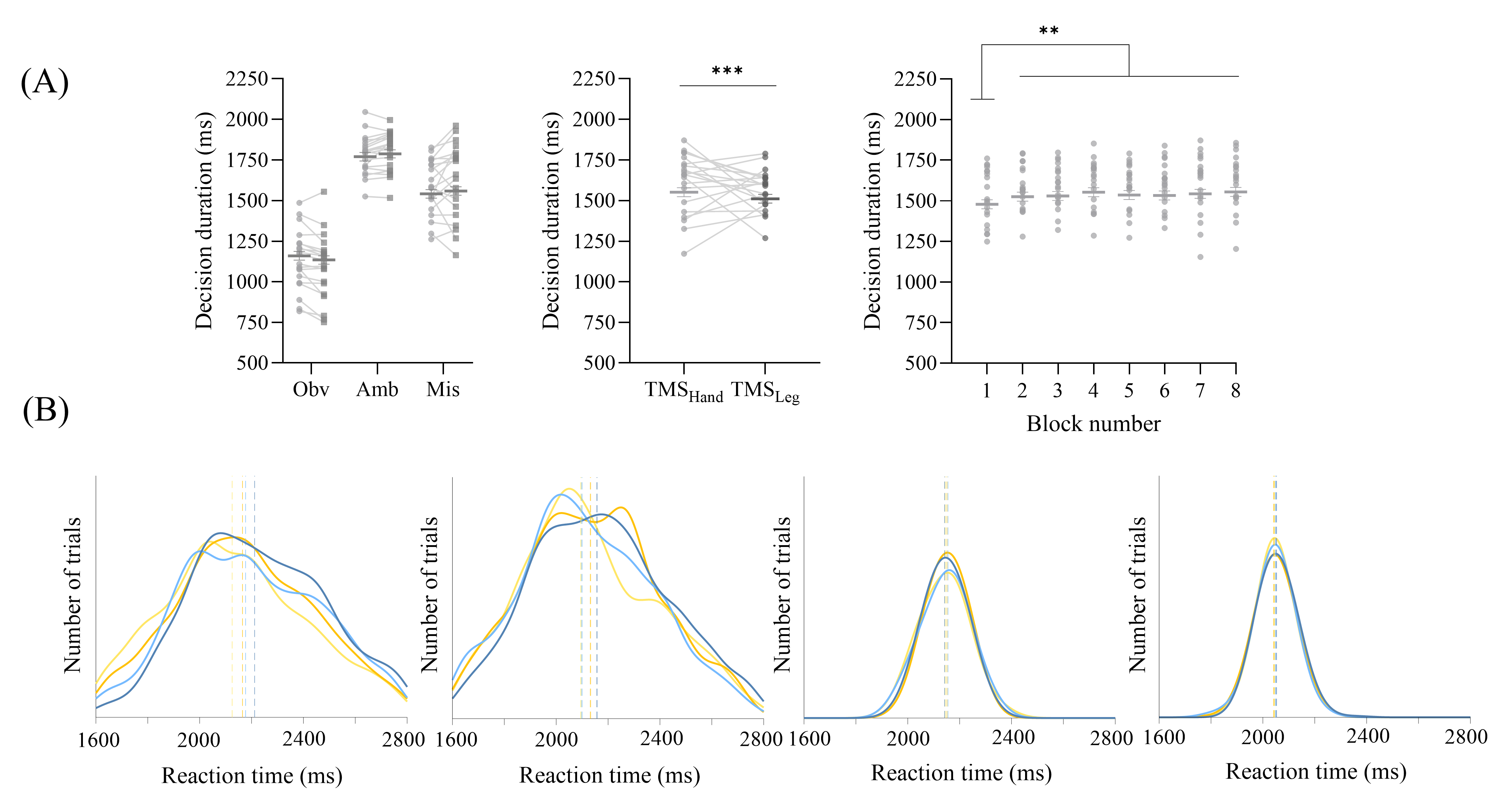

### Figure S2

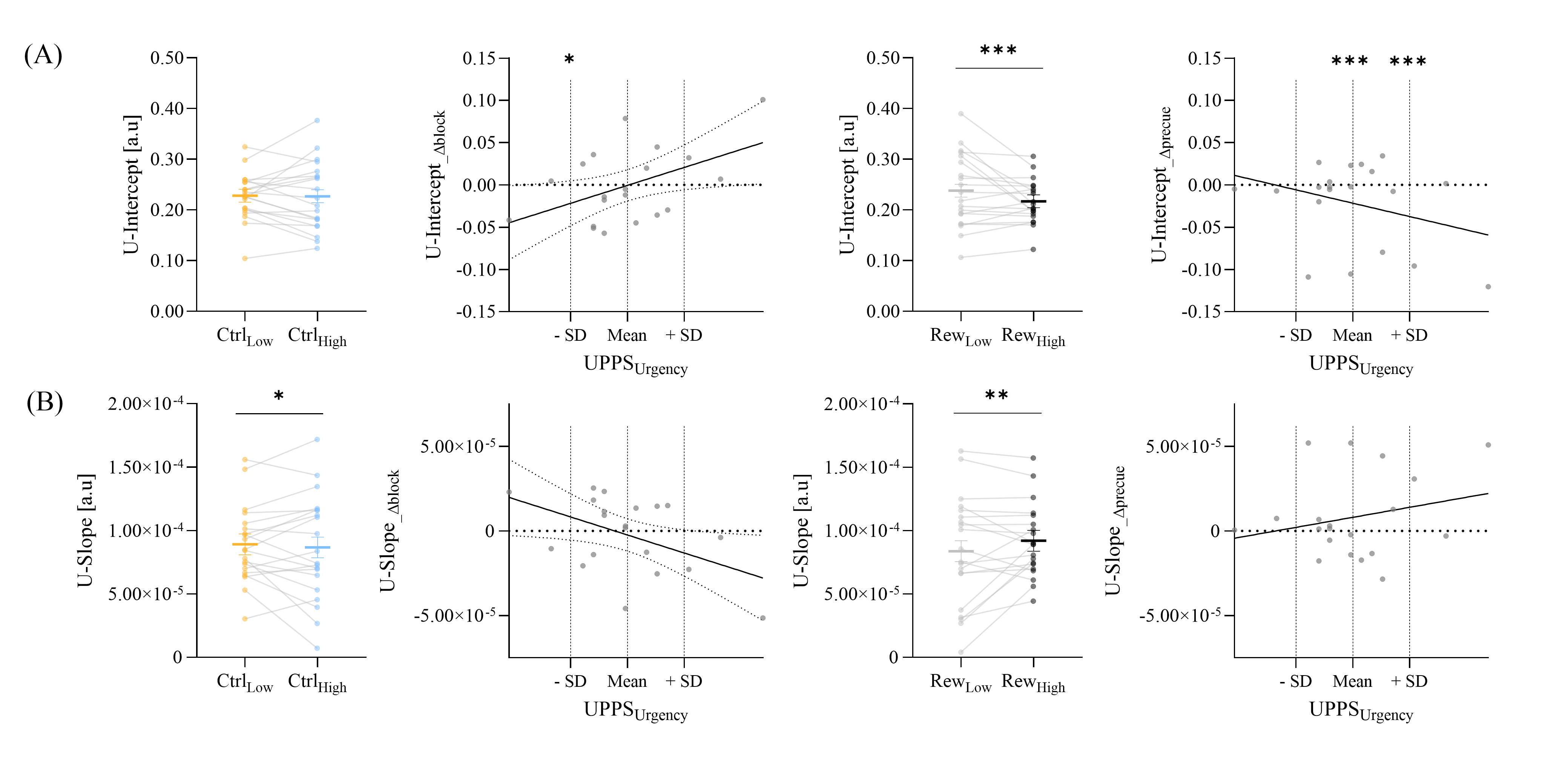

### Figure S3

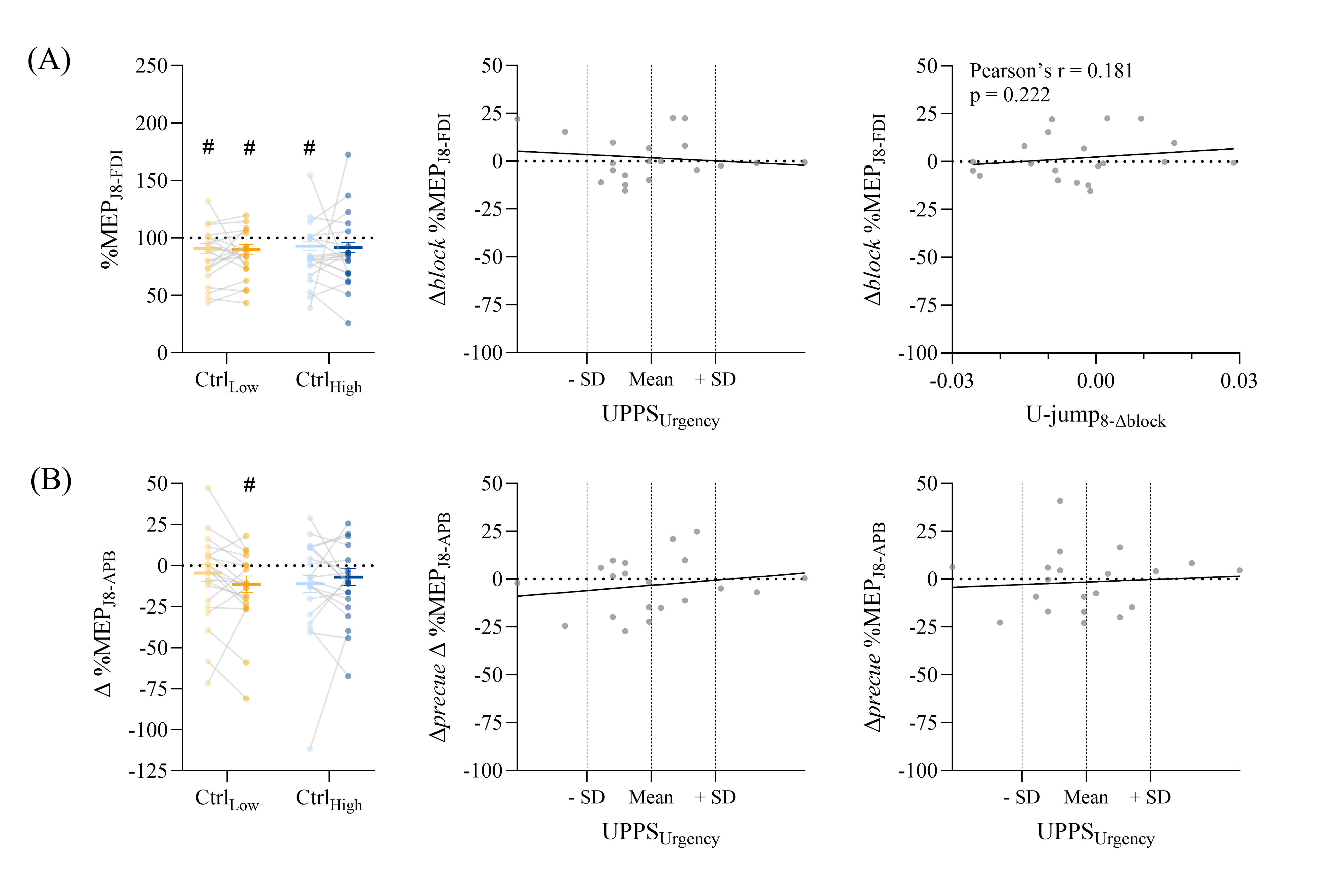
